# Distinct relations of microtubules and actin filaments with dendritic architecture

**DOI:** 10.1101/2019.12.22.885004

**Authors:** Sumit Nanda, Shatabdi Bhattacharjee, Daniel N. Cox, Giorgio A. Ascoli

## Abstract

Microtubules and F-actin have long been recognized as key regulators of dendritic morphology. Nevertheless, precisely ascertaining their distinct influences on dendritic trees have been hampered until now by the lack of direct, arbor-wide cytoskeletal quantification. We pair live confocal imaging of fluorescently labeled dendritic arborization (da) neurons in Drosophila larvae with complete multi-signal neural tracing to separately measure microtubules and F-actin. We demonstrate that dendritic arbor length is highly interrelated with local microtubule quantity, whereas local F-actin enrichment is associated with dendritic branching. Computational simulation of arbor structure solely constrained by experimentally observed subcellular distributions of these cytoskeletal components generated synthetic morphological and molecular patterns statistically equivalent to those of real da neurons, corroborating the efficacy of local microtubule and F-actin in describing dendritic architecture. The analysis and modeling outcomes hold true for the simplest (Class I), most complex (Class IV), and genetically altered (Formin3 overexpression) da neuron types.

SUPPORT: NIH R01 NS39600 and NS086082 and BICCN U01 MH114829.

## Introduction

Nervous systems comprise numerous neuronal subtypes, each with its own axonal-dendritic morphology and field of innervation. The specification and dynamic modification of subtype specific dendritic architecture not only dictate how distinct neuron subtypes form functional connections with other neurons, but also directly influences their computational properties (Lazarewicz et al., 2002; Ledda and Paratcha, 2017). The broad significance of dendritic form in neural function is further underscored by the wide spectrum of neurological and neurocognitive disorders that have been linked to disruptions in these processes (Franker and Hoogenraad, 2013). A vital challenge in understanding neuronal architecture is to elucidate how dendritic branches are regulated to elongate, bifurcate and terminate, leading to their unique tree geometry, and how these regulatory rules vary to yield the striking observed diversity in structure, function, and connectivity (Lefebvre et al., 2015). *Drosophila* larval sensory dendritic arborization (da) neurons have proven to be an exceptionally powerful model for investigating these questions, revealing mechanistic underpinnings of numerous structural dynamics including tiling, remodeling, outgrowth and subtype-specific branching (Jan and Jan, 2010; Nanda et al., 2017).

A combination of intrinsic genetic programming, extrinsic cues, electrophysiological activity and proximity to other cells (neuronal and non-neuronal) sculpt mature dendritic shape and maintain arbor stability as well as plasticity (Lefebvre et al., 2015). Each of these regulatory processes has been demonstrated to impact dendritic architecture by ultimately converging on the cytoskeleton (Franker and Hoogenraad, 2013; Jan and Jan, 2010; Ledda and Paratcha, 2017; Lefebvre et al., 2015; Nanda et al., 2017). Furthermore, the acquisition, maintenance, and modulation of diverse dendritic morphologies is largely mediated by organization and dynamics of F-actin and microtubule cytoskeletal components that provide the underlying scaffold and fiber tracks for intracellular trafficking critical to arbor development (Coles and Bradke, 2015; Franker and Hoogenraad, 2013; Kapitein et al., 2010; Kapitein and Hoogenraad, 2015). Numerous studies have demonstrated that diverse dendritic morphologies emerge via complex growth mechanisms modulated by intrinsic signaling involving combinatorial transcription factor regulatory codes that contribute to both neuronal subtype identity and morphological diversity (Corty et al., 2016; Das et al., 2017a; de la Torre-Ubieta L. and Bonni, 2011; Franker and Hoogenraad, 2013; Grueber et al., 2003; E. P. R. Iyer et al., 2013; S. C. Iyer et al., 2013; Li et al., 2004; Santiago and Bashaw, 2014; Sugimura et al., 2004; Ye et al., 2011). These transcription factor-mediated programs converge on a diverse set of cellular pathways to drive dendritic arbor diversity, many of which ultimately impact cytoskeletal architecture as a terminal mediator of arbor shape (Das et al., 2017a, 2017b; Ferreira et al., 2014; Franker and Hoogenraad, 2013; Hattori et al., 2013; Iyer et al., 2012; Jinushi-Nakao et al., 2007; Nagel et al., 2012).

Neuronal microtubules form a quasi-continuous core within the dendritic tree (Bray and Bunge, 1981). Microtubules result from the polymerization of α- and β-tubulin heterodimers on a structural template of γ-tubulin. The polarized protofilaments are then laterally arranged into a hollow configuration. Microtubules exhibit dynamic instability, *i.e*. an instant alteration between tubulin assembly and disassembly (Howard and Hyman, 2009). This switching can be intrinsically controlled by GTP/GDP binding to the heterodimers, where the GTP-tubulin lattice promotes polymerization and GDP-tubulin lattice promotes disassembly (Alushin et al., 2014). Unlike axons, whose microtubule rapidly polymerizing (“plus”) ends are uniformly oriented in the anterograde direction (Heidemann et al., 1981), dendrites can have mixed microtubule polarity, and microtubules remain significantly dynamic in mature dendritic trees. Specifically, in the fruit fly system, polymerizing microtubules exist in both proximal and distal dendrites, with a dominance of retrograde polymerization in proximal and anterograde polymerization in the distal (nascent) branches. Growing neurons without microtubule organizing centers regulate dendritic microtubule polymerization locally through nucleation (Sánchez-Huertas et al., 2016). Neurite diameter is also influenced by microtubule stability and organization via Ankyrin2 regulation (Stephan et al., 2015).

F-actin is also a polarized structure, consisting of ATP-bound globular actin primarily polymerizing at the barbed end (Coles and Bradke, 2015). All motile cells, including neurons, ubiquitously employ F-actin polymerization to drive plasma membrane protrusion, and actin-filled post-synaptic filipodia are involved in synaptogenesis (Ritzenthaler et al., 2000). F-actin polymerization is rate-controlled by nucleators such as Arp2/3 complex, which binds to monomeric actin to facilitate polymerization (Konietzny et al., 2017; Sturner et al., 2019) and enables F-actin polymer branching (Smith et al., 2013), and the formin family, which acts downstream of Rho-GTPases to nucleate unbranched F-actin (Konietzny et al., 2017). In addition to their recognized role in actin nucleation and polymerization (Breitsprecher and Goode, 2013; Chesarone et al., 2010; Kawabata Galbraith and Kengaku, 2019), formins have been also implicated in microtubule stabilization through Formin Homology domain induced acetylation (Thurston et al., 2012). A single molecule, Formin3, was shown to simultaneously regulate the stability and organization of the microtubule and F-actin cytoskeletons, thereby impacting dendritic architecture (Das et al., 2017b). Moreover, inward actin waves facilitated by ADF/Cofilin allow microtubules to protrude, stabilizing initial branch growth (Flynn et al., 2012). Recent fluorescence nanoscopic observations also revealed a periodic, ring-like, membrane-attached actin-spectrin scaffold mostly compartmentalizing axonal membrane, but also found in dendrites (Unsain et al., 2018).

Although numerous upstream regulatory pathways have been discovered for microtubules and F-actin, their distinct roles in defining overall dendritic structure are still actively debated. Studies across various experimental systems have revealed contradictory effects of cytoskeletal dynamics such as polymerization and stabilization. For example, depolymerized microtubule has been shown to invade filipodia leading to the formation of new branches. On the other hand, reducing polymerized microtubule by destabilizing microtubule limits dendritic outgrowth (Georges et al., 2008). Microtubule stabilization has also been shown to yield branching complexity in newly formed dendrites (Yau et al., 2014). Moreover, retrograde microtubule polymerization away from bifurcations and terminals restricts overall branching (Yalgin et al., 2015). While actin polymerization has been shown to drive neurite outgrowth (Chia et al., 2016), contrasting experiments have indicated that actin waves at growth cones do not initiate elongation and neurite outgrowth can occur without F-actin at neurite tips (Mortal et al., 2017). Possibly stabilizing actin blobs are also found at branch points (Nithianandam and Chien, 2018) and nucleation events have been involved in this punctate expression of F-actin (Sturner et al., 2019). Since the local densities of microtubules and F-actin have never been quantified arbor-wide in large numbers of neurons, the multifarious relation between arbor architecture and cytoskeletal composition remains incompletely understood.

Here we deploy the recently developed multi-modal confocal image stack tracing of multi-fluorescent neurons (Nanda et al., 2018a) to reconstruct a representative sample of mature da neuron morphologies containing arbor-wide local quantifications of microtubule and F-actin concentrations. We test whether the distributions of microtubule and F-actin in mature arbors can provide topological and developmental insights on the emergent dendritic structure. First we devise novel analyses of these enhanced data uncovering a doubly dissociated effect of microtubules and F-actin on dendritic architecture: local microtubule quantity predicts downstream dendritic length (for this and subsequent non-trivial terms, see Glossary in Supplementary Information) and thereby overall arbor size; conversely, tree complexity is associated with changes in local F-actin quantity, as bifurcating regions are enriched in F-actin. We then show that local cytoskeletal composition is sufficient in constraining complete arbor morphology by designing a generative model that computationally simulates tree structure purely based on local microtubule and local F-actin concentrations measured from real neurons. The analytical observations as well as computer simulation results remain robust across functionally distinct da neuron subclasses and genetic alteration.

## Results

Class I and Class IV neurons represent the two extremes in larval da neuron dendritic morphology with respect to arbor size and complexity (Iyer et al., 2013). Class I neurons exhibit the simplest arbors and are the smallest of da neuron subclasses (Figure 1 A, B, C, D). Conversely, Class IV neurons are space-filling and constitute the most complex and largest da neurons (Figure 1 E, F, G, H). Here we focused on the Class I ddaE neuron subtype and Class IV ddaC subtype. Moreover, we considered genetic perturbation of cytoskeletal regulation by examining Class IV (ddaC) neurons overexpressing Formin3 (Form3OE), which also differ significantly from the corresponding wild type cells in both size and complexity (Figure 1 I, J, K, L). Formin3 overexpression was selected for genetic perturbation based on prior analyses (Das et al., 2017b) demonstrating a molecular role in differentially regulating dendritic F-actin and microtubule cytoskeletal organization. All three neuron groups were subjected to live, *in vivo* imaging using genetically-encoded, multi-fluorescent transgenic reporter strains to visualize F-actin and microtubule distributions (Das et al., 2017a). All neurons were imaged during the 3^rd^ instar larva stage, when the da neurons acquire their mature shapes. Cytoskeletal organization in control or mutant genetic backgrounds was assessed by using Class I- or Class IV-specific GAL4 to label respectively stable microtubules via the mCherry-tagged microtubule associated protein Jupiter (Cabernard and Doe, 2009; Das et al., 2017a; Weiner et al., 2016) (Figure 1 A, E, I, Figure S1 A, E, I, Figure S2) and F-actin by a GFP-tagged Moesin actin binding domain (Figure 1 C, G, K, Figure S1 C, G, K) which has been previously validated and extensively utilized as a marker of F-actin (Anderson et al., 2005; Das et al., 2017a; Dutta et al., 2002; Jinushi-Nakao et al., 2007; Lee et al., 2011; Nagel et al., 2012; Nithianandam and Chien, 2018). Basic morphological reconstructions of these images consisted of vectorized tree structures composed of numerous, uniformly sampled, connected compartments. Enhanced multi-signal reconstructions were then created from the image stacks starting from the basic reconstruction templates by quantifying microtubule (Figure 1 B, F, J, Figure S1 B, F, J) and F-actin (Figure 1 D, H, L, Figure S1 D, H, L) intensities in every compartment. Microtubule (MT) intensities displayed a similar tapering pattern in all three neuron groups (Figure 1 B, F, J, Figure S1 B, F, J), generally decreasing with path distance from soma (negative Pearson correlation R coefficient for local MT quantity against path distance in all three neuron groups: −0.28 for Class I, −0.36 for Class IV wild type, and −0.52 for Class IV Form3OE) and centrifugal branch order (Pearson coefficients for MT against centrifugal branch order: −0.22 for Class I, −0.37 for Class IV wild type, and −0.43 for Class IV Form3OE). F-actin (F-act) intensities demonstrate weaker tapering tendency in Class I and Class IV wild type (WT) neurons (Pearson coefficient for local F-act quantity against path distance: −0.21 for Class I, −0.14 for Class IV WT, and −0.57 for Class IV Form3OE; for F-act against centrifugal branch order: −0.15 for Class I, −0.09 for Class IV WT, and −0.48 for Class IV Form3OE), particularly in Class IV WT neurons that had several terminal branches with high F-act (Figure 1 D, H, L, Figure S1 D, H, L).

**Figure 1.**
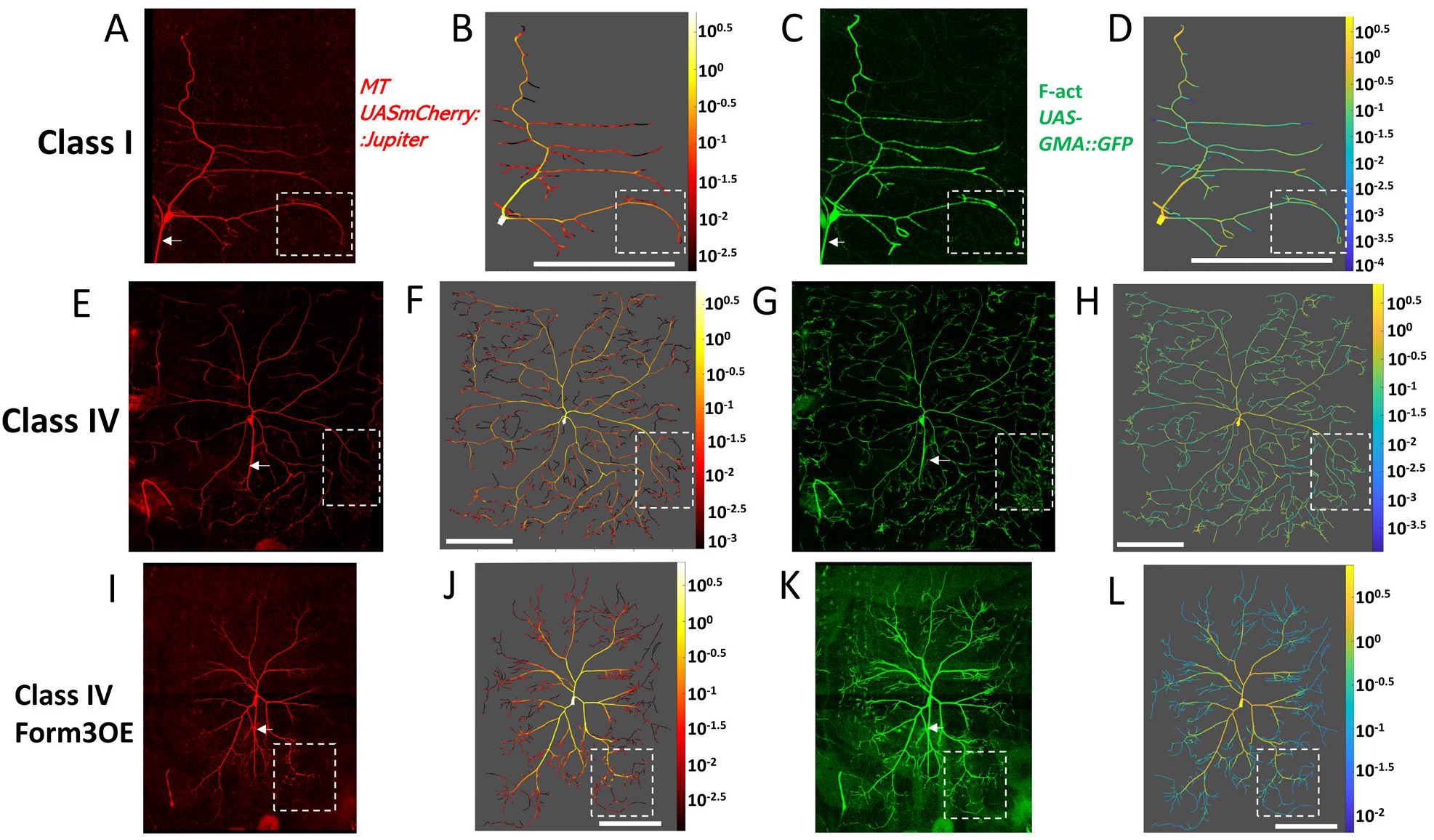
Confocal images and multi-signal reconstructions of Class I wild type (A, B, C, D), Class IV wild type (E, F, G, H) and Class IV mutant Form3OE (I, J, K, L). Microtubule (A, E, I) and F-actin (C, G, K) signals from confocal image stacks are traced together to produce multi-signal reconstructions, allowing the arbor-wide quantification and graphic rendering of microtubule (B, F, J) and F-actin (D, H, L) intensities (heatmaps represent cytoskeletal quantity). Axons (identified with arrows in A, C, E, G, I, K), only partially visible, are not traced and analyzed in this study. White scale bars (B, D, F, H, J, L): 100 μm. Figure S1 shows magnified versions of the areas demarcated by the dotted lines (A-L).

### Morphometric and cytoskeletal differentiation among neuron classes

As expected, quantitative morphometry demonstrated substantial differences among the three neuronal groups in overall size and morphology (Figure 2 A, D). Class IV WT neurons were clearly the largest in total dendritic length (19,305 ± 1510 μm, N=10) and had the largest spanning field (height 727± 63 μm, width 593± 32 μm). Genetically altered Class IV Form3OE neurons were smaller (total length 12,591 ± 985 μm, height 627± 44 μm, width 545± 32 μm, N=9), and Class I neurons were by far the smallest in length and spanning field (total length 1680 ± 166 μm, height 260± 21 μm, width 221± 21 μm, N=19). Class IV WT neurons also had the most complex dendritic arbors as measured by the number of branch terminals (683 ± 44.7), followed by Class IV Form3OE (365 ± 29.9) and the least complex Class I (26 ± 4.8). Correspondingly, the maximum Strahler (centripetal) orders for the three groups were 7, 6 and 4, respectively (Figure 2 D). These results are consistent with previous analyses of da neurons (Grueber et al., 2002; Nanda et al., 2018b).

**Figure 2.**
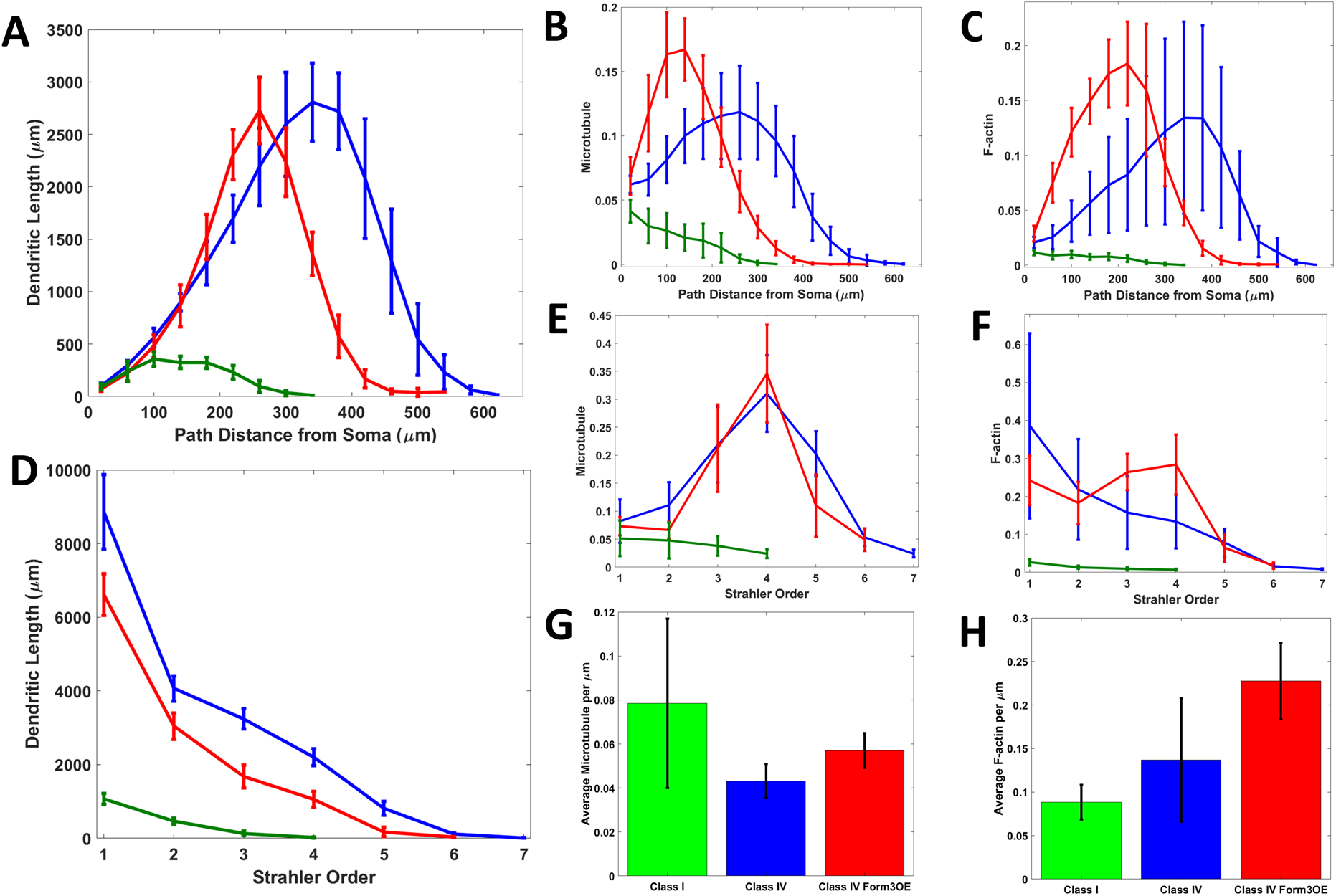
Morphological and cytoskeletal analysis of three groups of da neurons from the fruit fly larva. Distributions of dendritic length (A, D), microtubule (B, E), and F-actin (C, F) are examined as functions of path distance from soma (A, B, C) and Strahler order (D, E, F) for Class I (green), Class IV (blue) and mutant Class IV (red) neurons. Here dendritic length (in Y-axis) represents the total sum of dendrite length within each 40 μm path distance bin and at each Strahler order for every neuron, then averaged across neurons in each group. Similar to length, microtubule and F-actin quantities are summed for each path distance bin and Strahler order, then averaged across neurons within each group. In panels B, C, E, and F the Y axes were scaled so as to obtain unitary area under the curve for Class IV WT (blue lines), i.e. all cytoskeletal quantities in these four plots are represented relative to Class IV WT. Average microtubule (G) and F-actin (H) quantities are also reported per unit of length (μm), averaged across all neurons within a group. Error bars indicate standard deviation within group.

The morphological patterns were also reflected in overall intensity distributions for both MT (Figure 2 B, E) and F-act (Figure 2 C, F). Notably, on average, Class I had higher MT and lower F-act signal levels (per unit length) compared to Class IV WT and Class IV Form3OE, while Form3OE increased the average local quantities of both MT and F-act in Class IV neurons (Figure 2 G, H).

Interestingly, the local quantities of MT and F-act paralleled each other in the dendrites near the soma, but not farther out in the dendritic trees. Specifically, in the 20% most proximal dendrites, MT and F-act intensities were highly correlated (Pearson R: 0.98 for Class I, 0.88 for Class IV and 0.89 for Class IV Form3OE), whereas the 20% most distal locations displayed drastically reduced correlation (R= 0.46, 0.32 and 0.68 respectively for Class I, Class IV and Class IV Form3OE) (Table S1, Figure S3).

Tree asymmetry can be quantified as the average over all bifurcations of the normalized difference between the number of terminals (topological asymmetry) or the downstream arbor lengths (length asymmetry) stemming from each of the two children of a branch point. This metric ranges between 0 for a perfectly symmetric tree and 1 for a fully asymmetric tree (Van Pelt et al., 1992), and can often discriminate among neuron types or even between dendritic arbors within the same neurons, as in the apical and basal trees of pyramidal cells (Samsonovich and Ascoli, 2005). An alternative measure of dendritic asymmetry, caulescence, captures the degree to which a neuron has a main dendritic path or trunk (Brown et al., 2008) and weighs the imbalance of each bifurcation by the size of its sub-tree, thus emphasizing branches closer to the soma (see Methods for mathematical definitions of asymmetry and caulescence). While tree asymmetry did not vary greatly among the three groups of neurons considered here, there were substantial differences in tree caulescence. Class I neurons displayed the clearest main path as reflected by their topological caulescence (0.579 ± 0.091) and even more by length caulescence (0.631 ± 0.107). Class IV dendrites grow out in all directions, and hence did not show an evident main path (topological caulescence: 0.470 ± 0.080, length caulescence: 0.450 ± 0.067). Length and topological caulescence are significantly different between Class I and Class IV WT (p = 0.00005 and p = 0.0036), as well as between Class I and Class IV Form3OE (p = 0.0018 and p = 0.01) (Table S2).

### Local microtubule quantity is the best cytoskeletal predictor of arbor length

Total arbor length is a strongly discriminant descriptor of dendritic architecture and quantifies tree size by accounting for both overall span and branching complexity (Brown et al., 2008). Arbor length can be measured not only at the soma, where it captures the total dendritic elongation summed over the whole neuron, but also at any dendritic position considered as the root of the downstream sub-tree (Figure S4). Sub-tree arbor length is known to correlate with several geometric characteristics measured at the local root, including path distance from the soma, centrifugal branch order, and branch thickness (Donohue and Ascoli, 2005; Nanda et al., 2018b; Samsonovich and Ascoli, 2005). Therefore, these parameters have long been considered numerical proxies of underlying molecular processes that actually shape dendritic architecture (Burke et al., 1992; Donohue and Ascoli, 2008). The neuron-wide measurement of local MT and F-act quantities allowed us, for the first time, to test directly how the two major cytoskeletal mediators of dendritic growth compare with local geometric parameters in predicting downstream sub-tree length across the entire dendritic arbor. Path distance from the soma, centrifugal branch order, branch thickness, and downstream arbor length were calculated at every dendritic location in all neurons.

Among all local geometric and cytoskeletal molecular parameters analyzed, microtubule quantity had the highest correlation with arbor length (Pearson’s R coefficients, Table 1). MT quantity captured most arbor length variance (measured by the R-squared values) in Class IV Form3OE (77.4%) followed by Class IV WT (62.4%) and then Class I (44.9%). Higher quantities of MT co-occurred with greater arbor length and, when MT ran out, dendritic branches tended to terminate (Figure S4). Correspondingly, the correlation between MT quantity and arbor length was also considerably reduced in terminal branches (Table S1), where it only accounted for negligible amounts of variance (6.8% for Class I, 2.6% for Class IV and 0.04% for Class IV Form3OE). This is also consistent with the known greater dynamic plasticity of terminal branches, which can continuously elongate or retract in mature arbors. Interestingly, MT quantity alone had a higher correlation with arbor length than when combined with F-act quantity (Table 1). Among geometric parameters, branch diameter had the highest correlations with both MT quantity and arbor length (Table 1). Remarkably, these findings directly confirm early theoretical claims (Hillman, 1979; Burke et al., 1992). It should be noted that diameter measurement is limited by the resolution of light microscopy, which for the imaging system utilized in the current study is 311 nm (see Methods for calculation details). To estimate the maximum extent to which the correlation between arbor length and diameter could improve by eliminating the measurement error due to the resolution limit, we generated a surrogate set of diameter values by moving the measured values towards the regression line by a factor sampled from a uniform distribution with an upper bound of 311 nm (see Methods for details). Except for Class I, the resulting increased correlation still remained less predictive of arbor length than local MT (correlation between surrogate diameter and arbor length: 0.73 for Class I, 0.60 for Class IV, 0.67 for Form3OE). While this simulation provides an upper bound on the effect of the error, neurite thickness measured from electron microscopy will allow for a more accurate estimation.

**Table 1.** Pearson correlation coefficients (R-values) of arbor length against morphological and cytoskeletal parameters at the resolution of single (2 μm long) compartments. Microtubule quantity is the best predictor of arbor length for all analyzed cell groups.

| Cell Type | MT quantity | F-act quantity | MT + F-act quantity | Path distance from soma | Branch order | Diameter |
| --- | --- | --- | --- | --- | --- | --- |
| Class I | **0.67** | 0.55 | 0.62 | -0.47 | -0.33 | 0.64 |
| Class IV | **0.79** | 0.43 | 0.63 | -0.45 | -0.42 | 0.49 |
| Class IV Form3OE | **0.88** | 0.76 | 0.82 | -0.47 | -0.39 | 0.61 |

### Bifurcations are strongly associated with local F-actin enrichment

Relative to microtubules, F-actin quantity displayed weaker tapering tendency (except in Class IV Form3OE) and lower correlation with arbor length (Figures 1, 2, Table 1). Instead, we observed definitive patterns in the relative change of F-act between adjacent locations. Specifically, a systematic punctate expression of F-act was qualitatively apparent at dendritic bifurcations (Figure 3). To quantify this phenomenon, we computed the difference in F-act at every dendritic compartment relative to its parent compartment (ΔF), where a ΔF value of −1 represented complete elimination of F-act (from parent to the current), a ΔF value of 0 denoted no change, a ΔF value of +1 denoted doubling of F-act quantity (see Methods for detail). We then separated all compartments based on their topological behaviors: elongations, bifurcations, and terminations. We found that most bifurcating compartments (74% in Class I, 68% in Class IV, and 60% in Class IV Form3OE) showed F-act enrichment, *i.e*. an increase in F-act relative to their parent compartments. The cumulative ΔF frequency was significantly right-shifted for bifurcations relative to elongations and terminations in all three neuron groups (Figure 3, Figure S5). The median ΔF for bifurcations were 0.201, 0.121, and 0.041 in Class I, Class IV, and Class IV Form3OE neurons, respectively. The corresponding median ΔF for elongations were −0.024, −0.044, −0.033, respectively, indicating a tendency of F-act to decrease along non-bifurcating branches. Only 24% of Class I, 25% of Class IV and 31% of Class IV Form3OE bifurcations displayed a ΔF value lower than the respective elongation medians.

**Figure 3.**
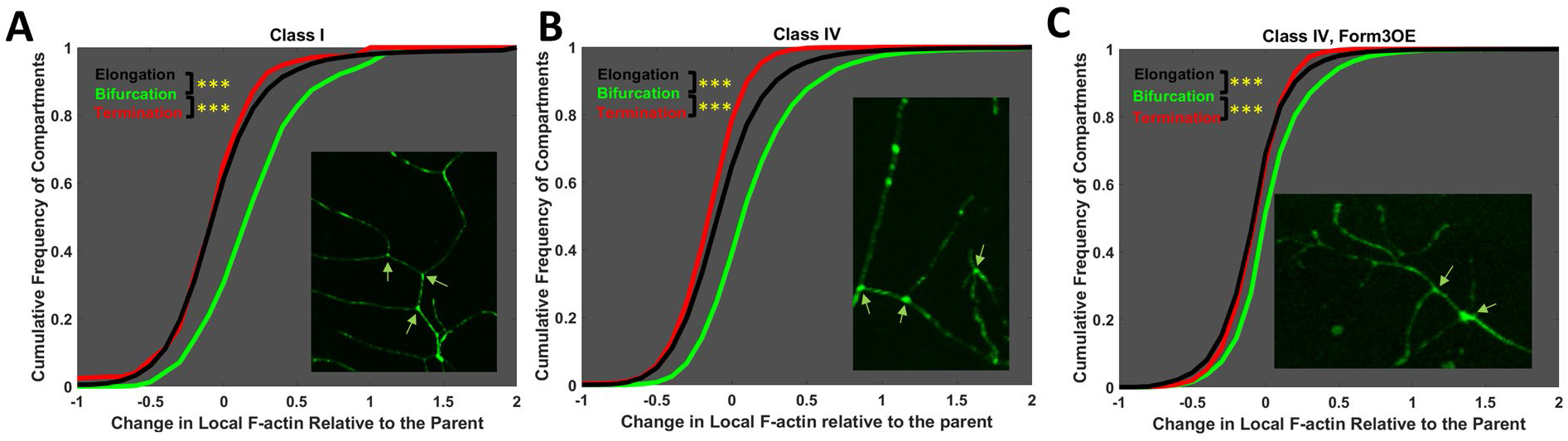
Cumulative frequency of the fold change in local F-actin concentration relative to the parent compartment for bifurcating (green), elongating (black), and terminating (red) compartments. A value of 0 indicates unchanged F-actin concentration; a value of −1 indicates complete F-actin disappearance (100% decrease relative to parent compartment); and a value of +1 indicates doubling F-actin concentration (100% increase relative to parent compartment). Class I (A), Class IV (B) and mutant Class IV Form3OE (C) distributions are shown separately. In all three cases, the bifurcating line is clearly right-shifted relative to the terminating and elongating lines. The arrowheads in the inset confocal images point to representative branch points. Triple asterisks indicate significantly different comparisons with p<0.001, Bonferroni-corrected 1-tail t-test.

To test whether F-act enrichment could be due to the enlargement of branch points, we repeated the analysis using signal intensity instead of F-act quantity, effectively normalizing the influence of dendritic diameter. This demonstrated equivalent or slightly more pronounced F-act enrichments, with median ΔF of 0.26, 0.15 and 0.07 at branch points in Class I, Class IV WT, and Class IV Form3OE neurons, respectively. Correspondingly, only 19% of Class I, 21% of Class IV WT, and 27% of Class IV Form3OE bifurcations displayed a ΔF intensity value lower than their respective elongation medians. Thus F-actin enrichment at bifurcations cannot be explained by branch point thickness. Furthermore, the levels of microtubule enrichments at branch points were modest when compared to those of F-actin. Class IV WT did not show any overall microtubule enrichment (median ΔM at branch points was 0). Class I showed relatively lower levels of MT enrichment (median ΔM of 0.101 at branch points, compared to ΔF of 0.201); Class IV Form3OE, which had the smallest levels of F-actin enrichment, had similar levels of MT enrichment (ΔM of 0.043 and ΔF of 0.041). These values can be explained simply by the correlation of microtubule and F-actin signals.

Recent in vivo studies have reported transient F-actin enrichment at da neuron branch points. One study demonstrated moving actin blobs that stabilize at branch points (Nithianandam and Chien, 2018). Another study showed that the Arp2/3 complex driven nucleation of actin initiates branch formation from the F-act enriched location (Sturner et al., 2019). To test whether F-act enrichment occurs preferentially or exclusively in newly formed branches, we analyzed enrichment as a function of Strahler order. Because all newly formed (terminal) branches have a Strahler order of 1, higher Strahler order branches must have formed earlier in the arbor development. We observed that more than half of all branch points across all Strahler orders, except the maximum order corresponding to the soma, were F-act enriched (Figure S6 A, B, C). Moreover, F-act enrichment moderately decreased with Strahler order in Class I (R = −0.17, P<0.05) and Class IV Form3OE (R = −0.11, P<0.05) neurons (Figure S6 D, E, F). While enrichment in higher Strahler orders suggests stable F-act enrichments in older branches, the greater level of enrichment in lower Strahler orders may reflect a dual mechanism of transient and stable enrichment in newly formed branches. The comprehensive arbor-wide quantification in the current study thus extends previous sparse observations of this phenomenon to mature arbors.

The average bifurcation angle (the local angle created at the branch point between the two bifurcating branches) was highest for Class I neurons (98 ± 6.1 degrees) followed by Class IV WT (92.8 ± 1.65 degrees) and Class IV Form3OE (90.09 ± 1.12 degrees). We tested if local bifurcation angle was correlated with MT, F-act or their local changes. While Class I WT neurons demonstrated no such correlations, Class IV WT neurons only showed very low negative correlation between local microtubule and bifurcation angle (R = −0.03, P<0.05). Class IV Form3OE neurons demonstrated low correlations between bifurcation angle and all four cytoskeletal parameters (Pearson R: MT = −0.08, F-act = −0.14, ΔM = 0.04, ΔF = 0.11).

### Distinct effects of microtubules and F-actin on dendritic growth and branching

Our quantitative analyses suggest a stark double dissociation between the two major mechanisms to establish dendritic architecture, namely branch elongation and bifurcation, and distinct cytoskeletal determinants, respectively microtubule quantity and F-actin enrichment. The smaller levels of residual association of branching with MT and arbor length with F-act can be attributed to the overall correlation between these two cytoskeletal quantities.

Correspondingly, an in-depth inspection of separate topological events (occurrence of terminating, elongating, or bifurcating dendrites, see Glossary in the Supplementary Information for definition) as a combined function of local MT quantity and relative F-act change (Figure S7 A, B, C) revealed almost entirely complementary cytoskeletal compositions underlying bifurcations (F-act enrichment independent of MT quantity) and terminations (strong decline or absence of MT). Only in Form3OE Class IV neurons did elongation-rich and termination-rich regions partially overlap (Figure S7 C). We also found converging evidence at the dendritic branch level. Class I neurons had a relatively high ratio of MT to F-act quantities (0.89 ± 0.32) and displayed greater average branch length (33.57 ± 5.11 μm), *i.e*. a reduced number of bifurcations relative to total elongation. Class IV neurons (both WT and Form3OE), on the contrary, had shorter branches on average (WT: 14.2 ± 1.0 μm, Form3OE: 17.3 ± 0.7 μm) and lower MT to F-act ratios (WT: 0.38 ± 0.14, Form3OE: 0.25 ± 0.04) (Table S3). In other words, the ratio between arbor size and complexity paralleled the ratio between microtubule and F-actin quantities. The proportion of arbor lacking detectable microtubule across neuronal groups was also negatively correlated with average branch length (R = −0.998, p <0.001) (Figure S8).

While F-act also correlates with arbor length and MT also tends to enrich at Class I and Class IV Form3OE branch points, both of these observations can be attributed to the correlation between MT and F-act. The differential associations of MT with arbor length and of F-act with branching is least specific in Class IV Form3OE neurons: in this group F-act correlates relatively strongly with arbor length (though not as much as MT) and the branch point enrichment of MT is equivalent to that of F-act. This could be explained by the higher correlation between local MT and F-act in Form3OE neurons (R= 0.86) as compared to Class IV WT neurons (R = 0.65).

Taken together, these interactions may provide a complete description of two fundamental aspects of dendritic arbor topology: size (total length) and complexity (number of branches). Considering this evidence, we proceeded to test whether local MT and F-act measurements could be sufficient to reproduce the arbor-wide morphological and cytoskeletal properties for all three neuron groups.

### Computational model of arbor generation based on local microtubule and F-actin

Previous studies have shown that neural arbors of various cell types (Ascoli et al., 2001; Donohue and Ascoli, 2008), including da neuron arbors (Nanda et al., 2018b), can be simulated by computational models constrained by local geometry (path distance, branch order, and branch thickness). Our analysis indicated that local cytoskeletal components had greater influence over arbor length and branching probability than previously used geometric parameters. To test the efficacy of local microtubule and F-actin quantities in reproducing overall tree architecture, we designed a computational model of dendritic morphology purely driven by local intensities of MT and F-act measured from the reconstructed dendritic trees (Figure 4). We directly tested whether local branching behaviors (extensions, branching etc.) can be statistically reproduced by two local cytoskeletal constraints, MT and F-act quantities, and whether the simulated structure match the topology of real neurons.

**Figure 4.**
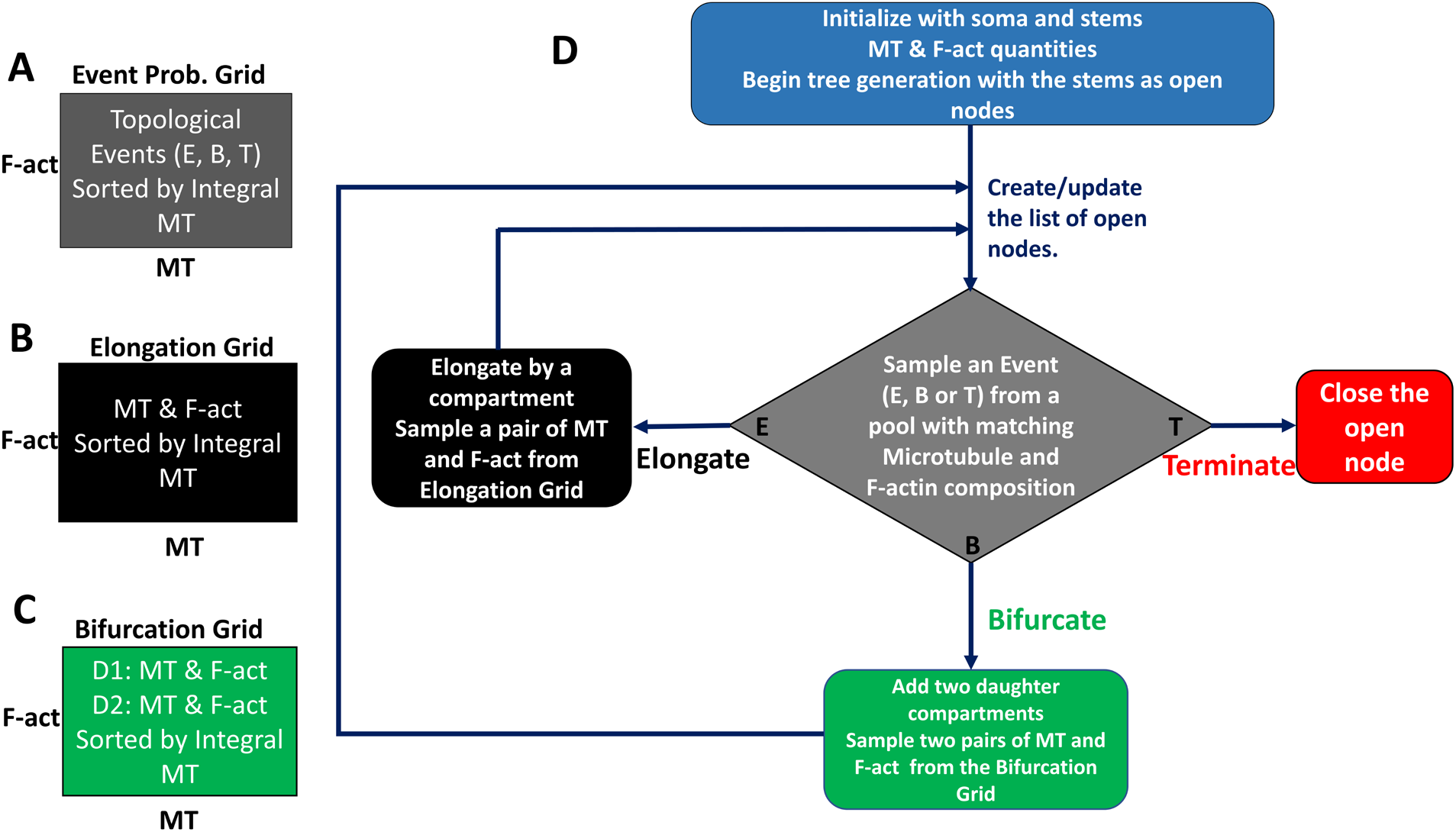
Generative model description. Stochastic sampling is carried out from three data grids (A, B, C) binned by local microtubule (MT) and F-actin (F-act) quantities. The grids are used to sample (A) branch topology (elongation, bifurcation or termination), as well as the cytoskeletal compositions for the child compartments of elongations (B) and bifurcations (C). All data are sampled from the bin that best matches the MT and F-act composition of the active compartment, with a bias for higher integral microtubule values (see Methods for definition). The tree generation process (D) starts with the initial MT and F-act concentration of the soma and the dendritic stem as active compartments. For each active compartment, branch topology sampling determines whether to elongate (adding one new compartment), bifurcate (adding two new compartments) or terminate (ending the branch). If a node elongates or bifurcates, the cytoskeletal composition of the newly created compartment(s) is sampled from the corresponding data grids. The process is repeated for all newly added nodes, until all branches terminate.

The model consisted of two stochastic processes iterated over all generated compartments starting from the dendritic roots stemming from the soma: sampling of topological events (elongation, bifurcation, and termination) and sampling of cytoskeletal composition. Thus, in these simulations, local MT and F-act quantities determined not only the branching behavior (whether to elongate, bifurcate or terminate) in that dendritic location, but also the cytoskeletal composition of the child compartment(s), which in turn determined the branching behavior of their child compartment(s), with the process continuing until all branches terminate. The first stochastic process sampled one of three possible topological events from the corresponding data grid (Figure 4A). Sampling an elongation event added one child compartment; sampling a bifurcation event added two child compartments; sampling a termination added no compartment and closed the active compartment. The second process, determining the cytoskeletal composition in the child compartments, was further subdivided into two sampling processes, for elongation (Figure 4B) and for bifurcation (Figure 4C). The elongation grid contained the cytoskeletal composition of all compartments whose parent were not bifurcating. The bifurcation grid contained the cytoskeletal composition of all bifurcation children. Depending on whether a compartment elongated or bifurcated, one (for elongations) or two (for bifurcations) pairs of MT and F-act quantities were sampled and assigned to the newly added dendritic compartment(s). All three grids (see Glossary in Supplementary Information) are two-dimensional matrices whose entries correspond to small ranges of MT and F-act values. The sampling processes were all similarly constrained by MT and F-act composition of the active compartment, ordered by the weighted microtubule integral from the soma to the compartment’s parent (see Methods for mathematical definition). The simulations ran until all compartments terminated (Figure 4D) and the number of simulated neurons was matched to the number of real neurons for every cell group. Note that this generative model only describes arbor topology and branch length (i.e., ‘dendrograms’) as well as cytoskeletal distributions, but not the three-dimensional geometry of dendritic trees (bifurcation angles and branch meandering). Both local parameters were essential to the model as high MT predicted larger downstream arbor length and enriched F-act separated bifurcation events from elongations and terminations.

### Simulated neurons reproduce morphology and cytoskeletal distributions across neuron groups

Dendrograms visually represent the size and complexity of neurons while disregarding other aspects of arbor shape such as angles. The x-axis of the dendrograms represent the number of bifurcations and terminations. The Y-axis represents path distance from the soma. Dendrograms are sorted so as to place the larger downstream subtree to the left of each bifurcation. The dendrograms generated by the computational simulations were visually and qualitatively equivalent (as if sampled from the same group) with those of the real neuron dendrograms for each of the three neuron groups (Figure 5).

**Figure 5.**
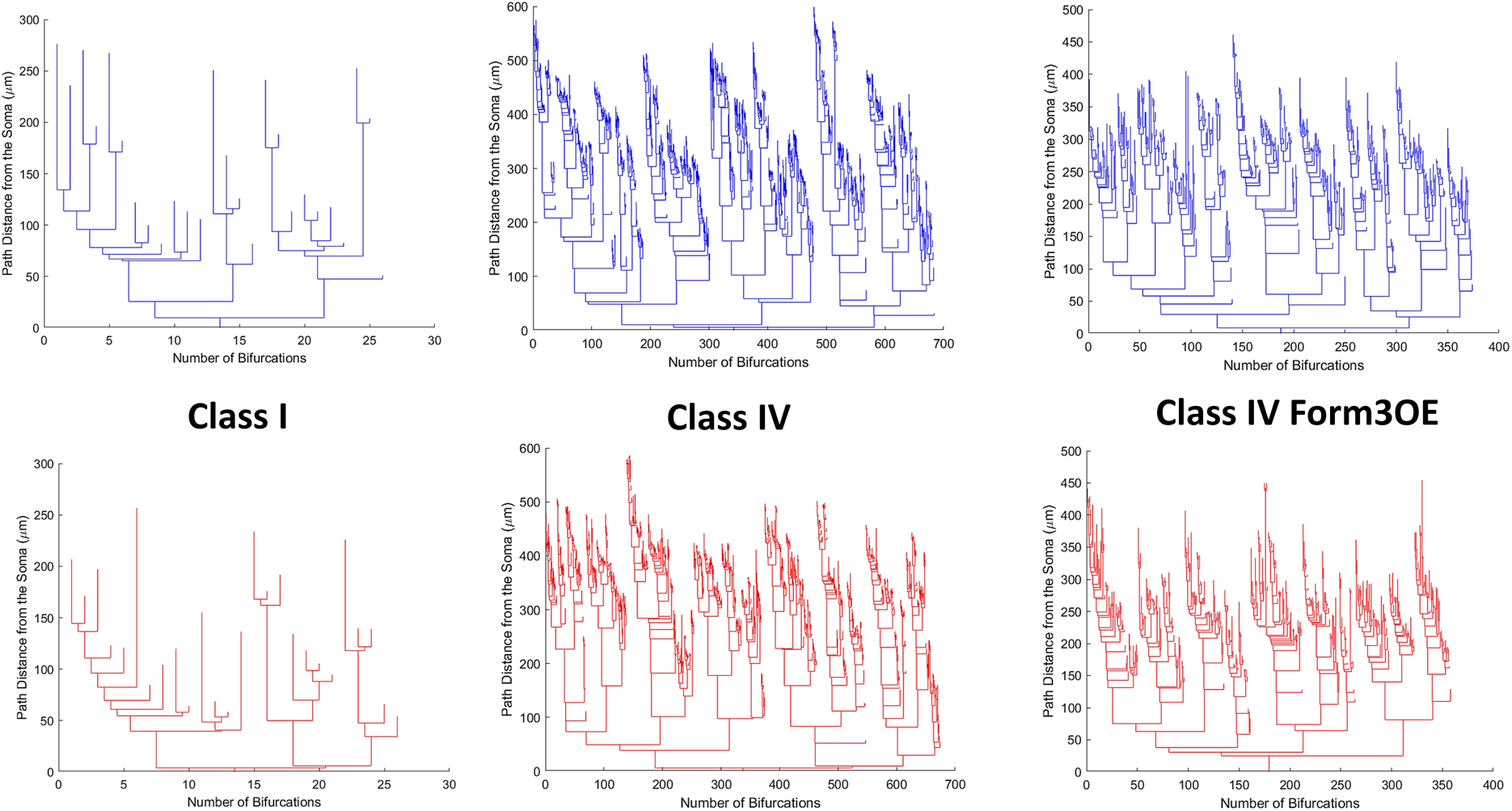
Representative dendrograms of real (top, blue) and simulated (bottom, red) Class I (left), WT Class IV (middle), and mutant Class IV Form3OE (right) neurons demonstrate the distinct size and complexity of the three neuron types. X-axis represents number of bifurcations and terminations; y-axis represents path distance from the soma. Dendrograms are sorted so that the larger subtree at each bifurcation is on the left.

Moreover, the simulated neurons of Class I, Class IV, and Class IV Form3OE quantitatively reproduced all measured morphometric characteristics of their real counterparts, namely total dendritic length, number of terminal tips, topological and length asymmetry, and topological and length caulescence. Thus, relative to the model-generated Class I dendrograms, model-generated Class IV dendrograms were much longer and more complex but had lower topological and length caulescence (Table S2). The model-generated Class IV Form3OE dendrograms had generally intermediate values between the two WT subclasses in these measures. In all quantified characteristics, the model fully reproduced the class-specific morphometrics in terms of group average and variance. None of the 18 two-tailed t-tests (6 measurements for each of 3 classes, Table S2) comparing real and simulated neurons yielded statistically significant differences (p > 0.5 even with no multiple testing correction).

Next, we compared real and simulated data by examining the distributions of dendritic length, microtubules, and F-actin as functions of path distance from soma and Strahler order. In all six cases the class-specific patterns of the simulated neurons were qualitatively and quantitatively almost identical to those of their real equivalents (Figure 6). Thus, the model was not only able to capture the overall tree properties of Class I, Class IV, and Class IV Form3OE, but also their rise or decline across path distance and branch order.

**Figure 6.**
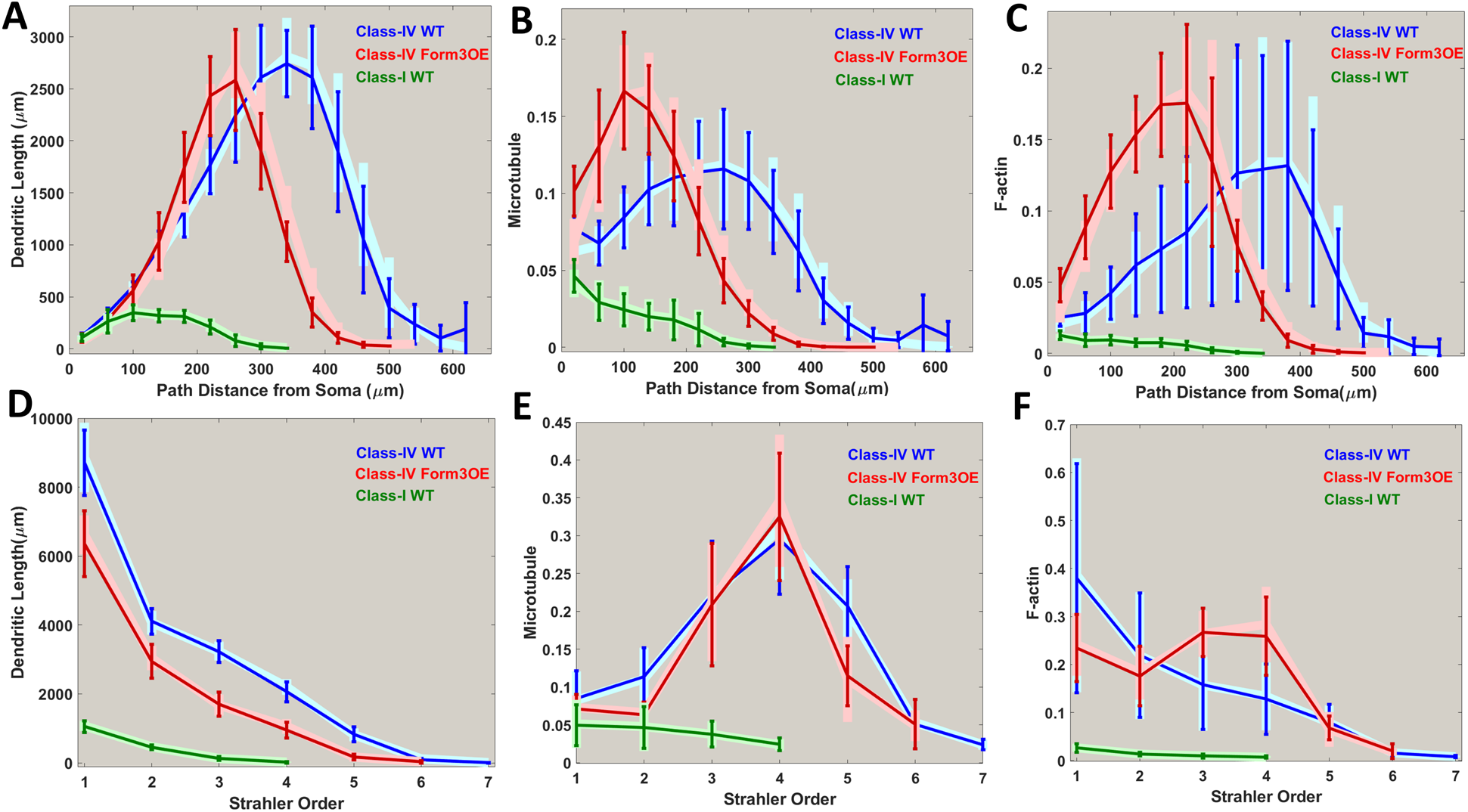
Emergent distributions of dendritic length (A, D), microtubule (B, E) and F-actin (C, F) by path distance (A, B, C) and Strahler order (D, E, F) of simulated Class I (green), Class IV (blue), and Class IV Form3OE (red) neurons. Error bars indicate standard deviation. Corresponding distributions of real neurons are overlaid in lighter shades of blue, red, and green.

Furthermore, simulated neurons exhibited similar relations between cytoskeletal composition and morphology as real neurons. Model-generated arbor length was highly correlated to local MT quantity (Table S4), but substantially less so in terminating branches (Table S1), and F-act enrichment was strongly associated with bifurcating compartments (Figure S9). Interestingly, the negligible correlation of arbor length with local MT in terminal dendritic branches did not significantly impact the similarity of real and simulated morphologies. This is because most of the terminal branches lacked detectable MT and, although the terminal branch lengths varied stochastically irrespective of MT quantity, such variability occurred within a relatively small range of arbor length when compared to overall arbor size.

Just as in real neurons, the correlation between MT and F-act quantities also decreased with path distance in synthetic neurons (Table S1, Figure S10). Finally, the computational model also reproduced the class-specific branch length (overall as well as separately for bifurcating and terminating branches) and the ratio between MT and F-act quantities (Table S3), thus replicating in the simulated neurons the parallel between the arbor size-to-complexity relation and the MT-to-F-act relation observed in real cells.

## Discussion

Microtubules and F-actin have long been implicated in neurostructural plasticity; however, their exact roles have not been fully elucidated due to the lack of their simultaneous quantification across the entirety of the neural arbor. With the use of the newly developed multi-signal reconstruction system (Nanda et al., 2018a), we quantified the arbor-wide microtubule and F-actin distributions from live images of fruit fly sensory neurons. Analysis of cytoskeletal distribution revealed the association of local microtubule with dendritic elongation (arbor size) and of local F-actin with dendritic branching (arbor complexity). These two distinct links of microtubules and F-actin with complementary aspects of dendritic morphology were consistent across structurally and functionally different cell types as well as a genetic alteration affecting cytoskeletal organization.

The striking structural difference in dendritic morphology between WT Class I and Class IV da neurons is reflected in their substantially different functional roles and efferent patterns within the larval sensory system. Class I neurons participate in proprioceptive functions (Cheng et al., 2010; He et al., 2019; Hughes and Thomas, 2007; Vaadia et al., 2019) whereas Class IV neurons are involved in polymodal nociception (Hwang et al., 2007; Lopez-Bellido et al., 2019; Tracey et al., 2003), chemosensation (Himmel et al., 2019), and photoreception (Xiang et al., 2010). Previous studies have also shown that genetic manipulations that alter overall morphology do not necessarily affect the size to complexity ratio of Class IV neurons. In contrast, Class I neurons drastically change their size to complexity ratio when genetically altered (Iyer et al., 2013). Despite these structural and functional differentiations, the architectural rules revealed in this study remained consistent across the two da class extremes, quantitatively relating local cytoskeletal composition to the immediate branching behavior (bifurcation, elongation or termination) on the one hand, and to overall morphology (arbor length and asymmetry) on the other. Intriguingly, class-specific dendritic architectures for either Class I or Class IV wild-type neurons could be computationally simulated with high fidelity by solely considering their different distribution patterns of local cytoskeletal compositions for F-actin and microtubules. Even genetic alterations in the mutant Class IV did not alter the fundamental relationships among local cytoskeletal composition, branching events, and overall arbor shape. The reduction in F-actin enrichment at branch points for Class IV Form3OE neurons relative to the WT was reflected in their relative reduction of overall branching complexity as well. We also observed across the three groups of neurons that the distribution patterns of microtubules were similar to those of F-actin in proximal, but not distal dendrites, which could be attributable to the constraints of transcription and transport: tubulin and actin monomers are produced at the soma, and then diffuse or are actively transported to the proximal branches first, followed by distal branches.

Most morphological characteristics analyzed in this study were statistically different between the three neuron groups. The proposed computational model, although simple, fully reproduced the group-by-group morphological and cytoskeletal properties observed in real neurons while demonstrating the correct relationships between cytoskeletal composition and arbor structure and vice versa. The morphological features captured by the model included total length and overall complexity, distribution of dendritic branches, asymmetry of length and branching, and prominence of main dendritic path. Cytoskeletal properties included arbor-wide distributions of microtubules and F-actin, correlation of microtubules with arbor length and association of branching with F-actin enrichment. Therefore, the analysis and modeling results jointly suggest that the diverse dendritic morphologies across all investigated cell groups shared similar underlying cytoskeletal mechanisms.

Consistent with the model, local increases in the quantity of stable microtubules through Formin3 overexpression further improved the correlation between microtubule quantity and arbor length. Increased F-actin signal by the same genetic alteration nearly doubled local F-actin quantity but actually reduced the enrichment levels at bifurcations. This suggests that enrichment events are more topology-specific in WT neurons, occurring when new branching events take place, while the tree-wide increase of F-actin in Form3OE decreases the difference between bifurcating and elongating compartments. As a combined result, Formin3 overexpression substantially lowered total dendritic length and arbor complexity in Class IV neurons, yielding a reduction in total F-actin and microtubule quantities despite the local increases of both cytoskeletal components. These observations corroborate the notion that Class IV neurons are evolutionarily optimized to fill a large amount of space through branching and elongation, maximizing the use of available resources (Grueber et al., 2002). Disrupting the distribution of cytoskeletal resources could thus interfere with its space filling mechanisms (Das et al., 2017a; Nanda et al., 2018b).

The consistent correlation observed between microtubule quantity and arbor length quantitatively confirms after forty years a seminal prediction based on early electron microscopy evidence (Hillman, 1979): namely, that the number of microtubule polymers progressively decreases along the dendrites with distance from the soma and approaches zero near the terminal tips. Specifically, this decrease was posited to be gradual within a branch, as some microtubules stop polymerizing, and discontinuous at bifurcations, where the microtubule polymers would partition between the two child dendrites. While the above conceptual model was originally formulated relying on sparse evidence from individual dendritic branches, it proved to remain consistent with more recent electron microscopic data demonstrating lack of microtubule in terminal branches in contrast to the microtubule “backbone” of primary dendrites (Schneider-Mizell et al., 2016). Our multi-signal dendritic reconstruction and analysis now quantitatively corroborates these earlier observations at the level of entire arbors and across neuron groups. Those pioneering ideas along with initial ultrastructural observations that dendritic branch thickness closely corresponds with the number of microtubule polymers at that location (Hillman et al., 1980) inspired a successful series of early computational simulations in which larger and smaller branch diameters increased the probability of bifurcating and terminating, respectively (Donohue and Ascoli, 2008; Schneider et al., 2012). Another resource-driven phenomenological computer model posited the diffusion or active transport of somatically-produced tubulin to the neurite tips or growth cones where it would polymerize to extend the existing microtubule structure. The dynamic competition of tubulin among outgrowing branches can simulate simple as well as complex arbor morphologies (Hjorth et al., 2014). Other computational models finessed the influence of anterograde microtubule movement in tree generation by reducing the likelihood of creating new branches with path distance from soma or branch order (Donohue and Ascoli, 2008; Nanda et al., 2018b; Samsonovich and Ascoli, 2005; Van Pelt and Uylings, 2002). The current study is unprecedented in driving the computational model directly by the experimentally recorded local cytoskeletal resources. While we observe that dendritic terminals mostly lacked visible microtubules, it is important to note that the microtubule marker Jupiter used in this study tags stable microtubules. Therefore, dynamic microtubules and tubulin monomers might be present in the terminal dendritic regions.

While local microtubule quantity predicted downstream arbor length, the local change in F-actin constrained branching events. Enrichment of F-actin near dendritic bifurcations also constitutes an arbor-wide validation of recent experimental observations. Actin blobs have been shown to stabilize at branch points, and nullifying that stabilization leads to a reduction in branch complexity (Nithianandam and Chien, 2018). Moreover, Arp2/3 complex temporarily localizes at dendritic bifurcations to nucleate actin, leading to actin remodeling at the nascent branch point (Sturner et al., 2019). Satellite Golgi outposts have also been observed at the dendritic branch points where microtubules are nucleated and membrane proteins are produced (Ori-McKenney et al., 2012). Higher F-actin levels may be important to maintain the presence of the Golgi apparatus at branch points since actin is involved in regulating Golgi structure and function (Gu et al., 2001). Interestingly, the differences in F-actin enrichment between terminating and elongating dendrites were much more modest. This may be because the reconstructions are from static image stacks, providing a momentary snapshot of a structurally dynamic arbor. Hence, the compartments annotated as terminating could either be elongating or retracting. This interpretation is consistent with previous experimental observations that terminal branches remain dynamic even when the primary branches become stable (Lee et al., 2011).

While the local cytoskeletal composition of mature arbors was surprisingly predictive of overall dendritic size and complexity, this study did not model the geometric shape of the arbors, which results from a combination of several factors including stem orientation (Samsonovich and Ascoli, 2003) and branch angles (Bielza et al., 2014), self-avoidance (Kidd and Condron, 2007; Sundararajan et al., 2019), and tiling (Grueber et al., 2002). Dendritic pruning (Herzmann et al., 2018) and underlying developmental processes such as the polymerization, depolymerization, bundling, severing of microtubules or F-actin, traffics of cargos, and stoichiometry of microtubules and F-actin are also not accounted for in the model. Time-lapse imaging of dendritic morphology with multiple fluorescent proteins tagging the orientation of microtubule and F-actin polymerization will be informative for future studies to investigate further arbor plasticity and the underlying subcellular changes that drive it. We have recently developed an enhanced reconstruction system to annotate temporal changes in neuronal structure simultaneously with subcellular changes (Nanda et al., 2018a). Time varying reconstructions with local polymerization information (rate and direction) may foster a deeper understanding of potential developmental mechanisms. Future studies should attempt to directly alter microtubule or actin stability by using cytoskeletal effectors such as drugs (Otto et al., 1979), increase in temperature (Spedden et al., 2013), and mechanical force to observe the effects on overall neuron morphology. The sufficiency of newly hypothesized rules of growth can then be tested by implementing a computational simulation constrained by the enhanced acquired data to verify whether it also reproduces dynamical phenomena like scaling, retraction, and branch thickening. However, in order to become viable for use in developmental models, time-lapse reconstructions require the careful registration of image stacks, which is still exceedingly challenging at the level of entire dendritic arbors *in vivo* (Rada et al., 2018; Sheintuch et al., 2017).

## Methods

### *Drosophila* strains and live confocal imaging

*Drosophila* stocks were reared at 25°C on standard cornmeal-molasses-agar media. Age-matched wandering third instar larvae of both sexes were used for all experiments. Fly strains used in this included: *w1118, UAS-GMA::GFP; GAL4[477],UAS-mCherry::Jupiter* (Das et al., 2017a) (Class IV cytoskeletal reporter strain); *w1118, UAS-GMA::GFP;+;GAL4[221],UAS-mCherry::Jupiter* (Class I cytoskeletal reporter strain); *UAS-form3-B1* (Tanaka et al., 2004). *w1118* was used as a genetic background control for outcrosses to the multi-fluorescent cytoskeletal transgene reports. We previously confirmed that expression of the F-actin and microtubule transgene reporters did not themselves exert any effects on dendritic development (Das et al., 2017a). Live confocal imaging of age-matched third instar larval mature da neuron Class I and Class IV arbors was performed as previously described (Das et al., 2017a; Nanda et al., 2018a, 2018b). Briefly, larvae were immersed in a 1:5 (v/v) diethyl ether to halocarbon oil on a cover slipped glass slide and neurons expressing fluorescent protein cytoskeletal reporter transgenes were visualized on a Zeiss LSM 780 confocal microscope. Dissection and immunofluorescent labeling of third instar larval filets was performed as previously described (Sulkowski et al., 2011). Primary antibodies used in the study were rabbit anti-mCherry (used at 1:500) (Abcam; catalog # ab167453) and mouse anti-Futsch (22C10; used at 1:200) (Developmental Studies Hybridoma Bank; catalog # 22C10) (See Figure S2 for comparison of MT signal between UAS-mCherry::Jupiter and Futsch). Secondary antibodies were donkey anti-mouse Alexa fluor 488 (1:200) (Invitrogen; catalog # A21202) and donkey anti-rabbit Alexa fluor 555 (1:200) (Invitrogen; catalog # A-31572). Images were collected as z-stacks at a step size of 1-2 microns and 1024 × 1024 resolution using a 20X air objective (Plan Apo M27, NA 0.8) and 1 airy unit for the pinhole. A total of 19 Class I ddaE, 10 Class IV ddaC, 9 mutant Class IV ddaC Form3OE neurons were imaged, reconstructed, analyzed, and modeled. The digital morphological tracings and enhanced multi-signal ESWC files of all 38 neurons have been deposited to NeuroMorpho.Org (Ascoli et al., 2007) as part of the Ascoli and Cox archives. Images and the processed data have been deposited in Mendeley.com along with the analysis and modeling code (DOI: doi.org/10.17632/v3ncxmj6fn.1) for open access release upon publication of this manuscript.

### Neuromorphometric analyses

Confocal images were semi-automatically reconstructed and edited in neuTube (Feng et al., 2015). Larval da neurons are relatively flat with limited dendritic arborization in the Z dimension. Therefore, images containing high levels of noise (usually the top and bottom of the Z stacks) were removed and maximum intensity projection of the remaining stacks were used to guide the reconstructions process. The MT signal from the red channel and F-act signal from the green channel were combined in FIJI (Schindelin et al., 2012) to create a third pseudo membrane channel from which the initial tracing was carried out. TREES Toolbox (Cuntz et al., 2010) was then used to resolve inaccuracies in tree topology and to resample the dendritic tree into compartment of uniform length (approximately 2 μm). These edited and resampled trees where then manually curated in neuTube once more and scaled from pixels to physical unit (μm) to generate the final SWC files. To test whether the morphologies imaged from the combined MT and F-act signal used in this study match with previous membrane tagged tracings, we downloaded legacy Class IV ddaC (N= 16) and Class I ddaE (N= 19) neural reconstructions from NeuroMorpho.Org (Nanda et al., 2018b; Sulkowski et al., 2011) and compared their overall size and complexity with the neurons from the current study. The total dendritic lengths were not statistically different (P>0.05, two-tailed t-test) in either Class IV (18,753±5704 μm from NeuroMorpho.Org vs. 19,305±1510 μm in the current study) or Class I neurons (1897±489 μm from NeuroMorpho.Org vs. 1680±166 μm in the current study).

Multi-signal ESWC reconstructions (Nanda et al., 2018a) were produced in Vaa3D, using the *Multichannel_Compute* plugin, with the reconstructed SWC files and the multichannel images as input and a threshold of 10 for both channels (i.e. any intensity below 10 out of 255 was considered as 0). Specifically, the plugin was run twice, first with the basic SWC as the reconstruction and MT (red) as the secondary channel; and then a second time with the resultant multi-signal SWC from the first run as the reconstruction and F-act (green) as the secondary channel. A combination of the red and green channels was used as the primary channel for both runs. The final multi-signal reconstructions contained sub-structural information for both MT and F-act. The two signals were quantified in each compartment as the sum over all pixels in that compartment of the corresponding (MT or F-act) intensity divided by the maximum value of 255 to renormalize between 0 to 1 and reported per unit of length.

Custom scripts were written leveraging the TREES Toolbox (Cuntz et al., 2010) functions to calculate and plot multiple topological and cytoskeletal properties both locally and as arbor-wide distributions, including analysis of dendritic length and cytoskeletal protein expression vs. path distance from soma and Strahler order. Cytoskeletal quantities were normalized based on average total MT and average total F-act of Class IV WT neurons. We analyzed a large number of dendritic compartments from all three neuron groups: 16,052 from Class I (15,079 elongations, 477 bifurcations and 496 terminations), 98,520 from Class IV WT (84,872 elongations, 6819 bifurcations, 6829 terminations), and 66,305 from Class IV Form3OE (59,738 elongations, 3279 bifurcations, 3288 terminations).

### Mathematical definitions of asymmetry and caulescence

Topological asymmetry (Van Pelt et al., 1992) is defined as the average over all neuron bifurcations of the following function:

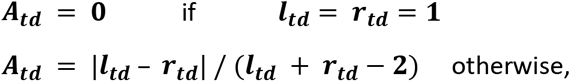

where ***l_td_*** and ***r_td_*** represent the numbers of terminal tips (or terminal degree) respectively in the left the right subtrees of the bifurcation.

Length asymmetry is similarly defined as the average over all neuron bifurcations of the following function:

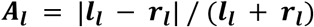

where ***l_l_*** and ***r_l_*** represent the arbor length respectively of the left the right subtrees of the bifurcation.

Topological caulescence (Brown et al., 2008) is the weighted topological asymmetry across the dendritic path from the soma to the terminal with the highest centrifugal branch order:

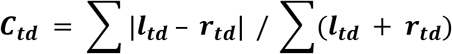

Length caulescence is the weighted length asymmetry across the dendritic path from the soma to the terminal with the highest path distance from the soma:

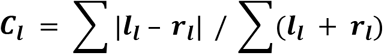

### Estimation of image resolution and its influence on dendritic diameter correlations

We calculated the theoretical resolution limit of the imaging system utilized in this study using the full width half maximum (FWHM) definition, whereas the lateral resolution in confocal laser scanning microscopy is given by 0.51*lambda/NA, where lambda is the excitation wavelength and NA is the numerical aperture. In our case, NA = 0.8 and lambda = 488 (corresponding to GFP). Using these formula and parameters, we calculate a lateral resolution of 311.1 nm. Since the imaging system is unable to differentiate objects below this resolution, we use 311.1 nm as the maximum error value to estimate the theoretically greatest possible increase in linear correlation between diameter and arbor length. We treat the difference between the real (measured) and the expected (based on its downstream arbor length) diameter value as noise and proceed to reduce it by a correction factor sampled for each neuronal compartment from a uniform distribution bounded by the smaller of the resolution limit (311.1 nm) and the measured difference for that compartment. We run the simulation 10 times for each neuron type and calculate the average correlation across the 10 trials. While, as expected, correlation increases for all neuron groups, they remain less predictive than local microtubule except for Class I (see Results).

### Relative change in F-actin

We first calculated the change in F-act quantity of every compartment from its parent compartment by simply subtracting the current quantity of F-act from that of the parent compartment. We then compute the ratio of F-act change to the parent F-act quantity to yield the relative F-act change (FΔ):

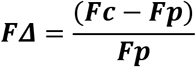

where Fp and Fc are the F-actin quantities of the parent and of the current compartment, respectively.

### Microtubule and F-actin based tree generation model

The data-driven model to simulate synthetic dendritic arbors was implemented in MATLAB (MathWorks, Natick, MA). This stochastic model uses three sampling grids: one for topological events and two for cytoskeletal composition (one each for elongating and bifurcating compartments). These grids can be thought of as two-dimensional matrices whose individual entries are populated with appropriate data extracted from real neurons (topological event in the first grid, cytoskeletal composition in the second and third) binned by the corresponding local microtubule and F-actin quantities.

The simulations started by sourcing the dendritic roots stemming from the soma of a real neuron as the ‘active’ compartments. Then, for every active compartment, the model identified the topological grid bin most closely matching the microtubule and F-actin quantities and sampled one of three topological events (elongation, bifurcation or termination) according to the following process. Data in the bin were first sorted by descending integral microtubule (***IM***), defined for the (i+1)^th^ compartment as:

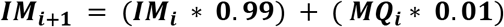

where ***MQ*** is the microtubule quantity and the i^th^ compartment is the parent of the (i+1)^th^ compartment. Integral microtubule can equivalently be expressed as:

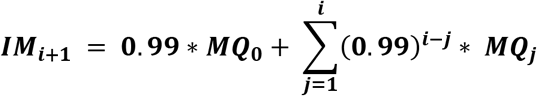

where ***MQ_0_*** is the microtubule quantity at the root compartment. The integral microtubule quantifies the progression of microtubule across the dendritic path from the soma to the current location. For sampling within a bin of n elements, first a number m was randomly chosen between 1 and n. This was followed by choosing a number between 1 and m and the process was repeated four additional times (totaling six iterations). This procedure reduces the effect of noise in the real data and increases the probability of sampling data points with higher integral microtubule earlier during a simulation. Once an element was sampled, it was removed from the data bin. If an elongation was sampled, a new compartment was added, sampling its cytoskeletal composition from the elongation grid with the same process described above. In case of bifurcations, two new compartments were added sampling their cytoskeletal compositions from the bifurcation grid. In case of termination, the compartment ‘inactivated’, ending the branch. The simulations stopped when all compartments inactivated.

All analysis and modeling code have been deposited to Mendeley.com (DOI: http://dx.doi.org/10.17632/v3ncxmj6fn.1) for open access release upon publication of this manuscript.

## Author Contributions

S.N. and G.A.A. originally designed the paper and conceived the model with D.N.C. providing early ideas on experimental data. S.B. conducted all wet lab experiments and multi-signal neuronal image acquisition with D.N.C’s supervision. S.N. reconstructed the neurons and wrote all analysis and simulation software. S.N. and G.A.A. analyzed the data and devised the generative algorithm of dendritic arbor morphology. S.N. created the figures and tables by incorporating feedback from all co-authors. S.N. and G.A.A. wrote the manuscript with D.N.C. and S.B. providing crucial feedback and edits. D.N.C. and G.A.A. are responsible for funding acquisition and project management.

## Acknowledgements

The authors thank Mahan Mollajafar, Anna Lulushi, Akhil Goel, and Alisha Compton for helping with neuron tracings.

