## Supplementary figures and tables for "Distinct relations of microtubules and actin filaments with dendritic architecture"

### Supplementary Information:

Link to data and code (Mendeley.Com):

DOI: http://dx.doi.org/10.17632/v3ncxmj6fn.1

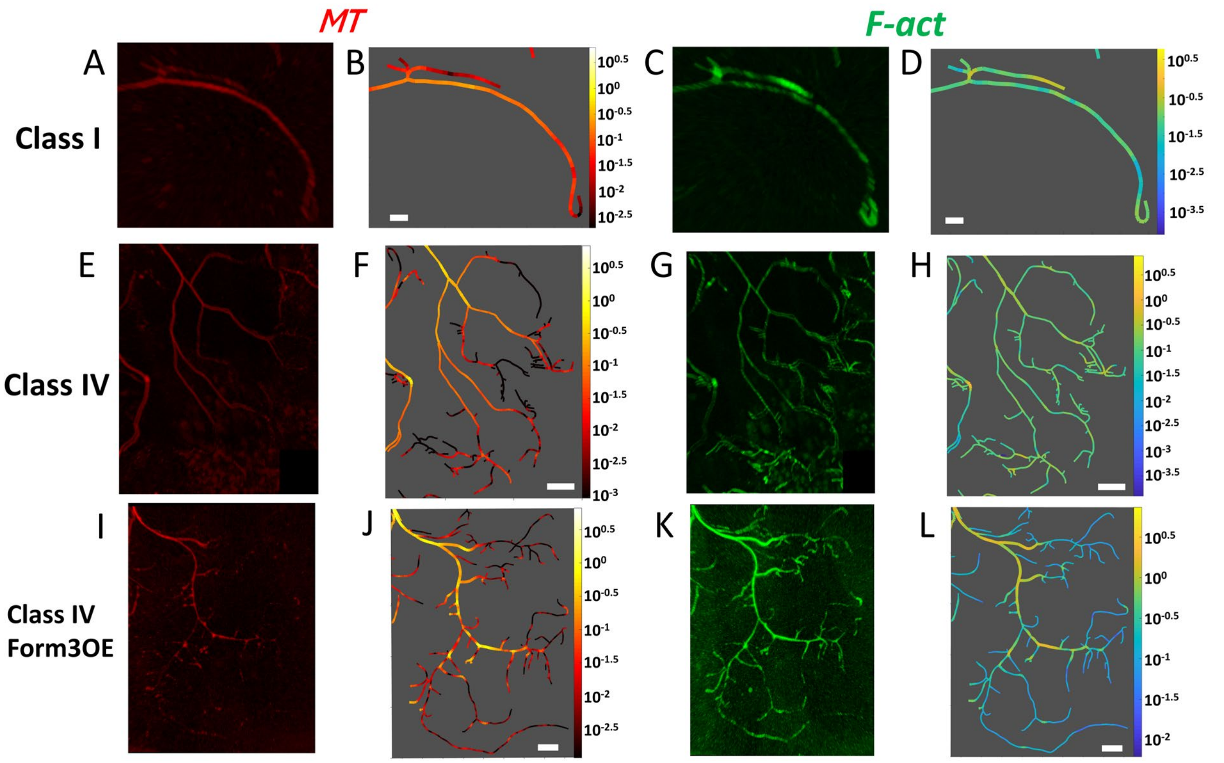


***Figure S1*** *Magnified version of the demarcated regions from Figure 1. Confocal images and multi-signal reconstructions of Class I wild type (A, B, C, D), Class IV wild type (E, F, G, H) and Class IV mutant Form3OE (I, J, K, L). Microtubule (A, E, I) and F-actin (C, G, K) signals from confocal image stacks are traced together to produce multi-signal reconstructions, allowing the arbor-wide quantification and graphic rendering of microtubule (B, F, J) and F-actin (D, H, L) intensities. Here we observe three terminal subtrees from Class I, Class IV and Class IV Form3OE neurons.*

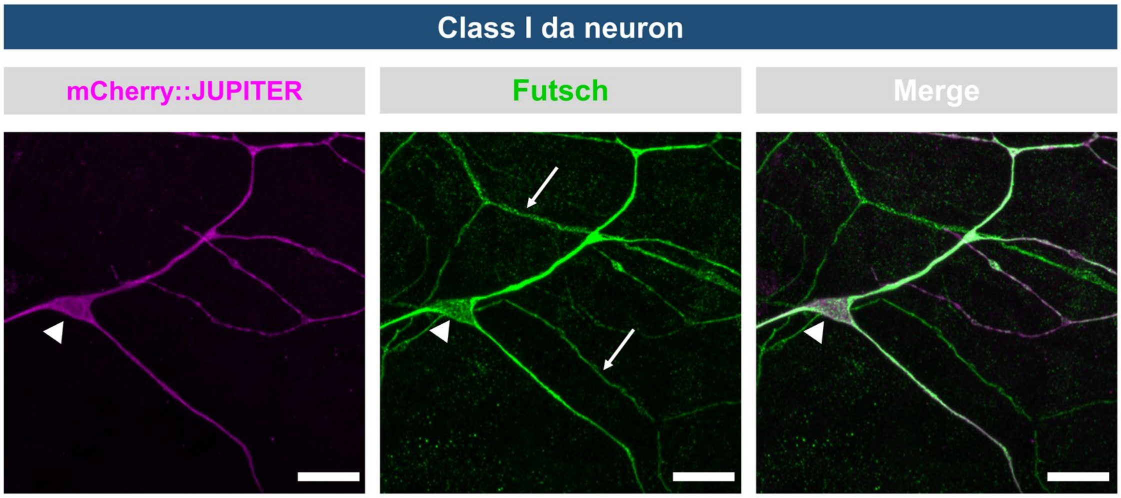


***Figure S2*** *Representative image of third instar larval Class I (ddaE) neuron labeled by class-specific GAL4[221] driven expression of UAS-mCherry::Jupiter to mark stable microtubules (magenta) (n=10 class I neurons). Larval filets were subjected to immunohistochemical labeling with antibodies against the microtubule binding protein Futsch (green), which labels all dendrites of all da neuron dendritic arbors (classes I-IV), and mCherry to mark the expression of the microtubule binding protein Jupiter (magenta). The location of the Class I ddaE cell body is denoted by the arrowhead in each panel, while the arrows mark dendrites emanating from other adjacent da neuron subtypes that are not labeled by the Class I driven Jupiter expression. The merge image clearly reveals overlap between these two microtubule binding proteins. Scale bar corresponds to 10 microns.*

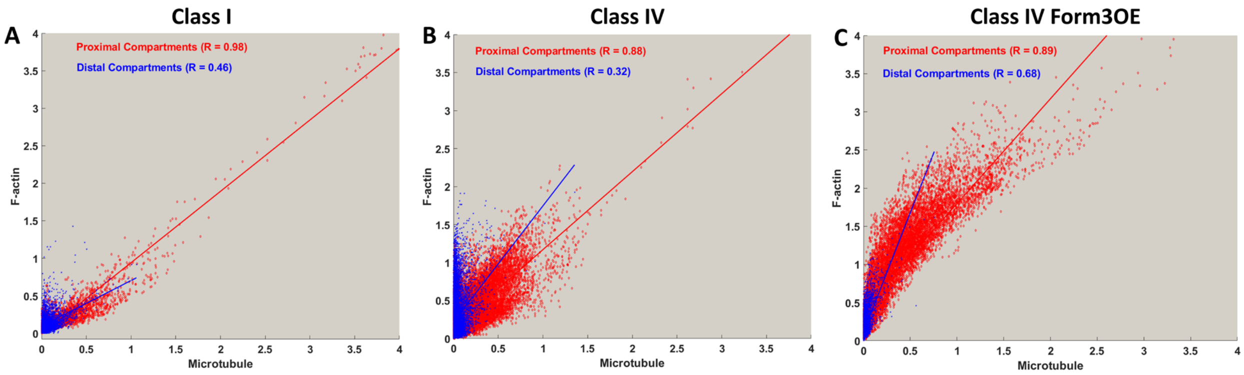


***Figure S3*** *The distribution of F-actin as a function of microtubule in proximal (the closest 20% of the compartments, in Red) and distal (the farthest 20% of the compartments) dendritic compartments from Class I (A), Class IV WT (B) and Class IV Form3OE (C) neurons. The correlation between microtubule and F-actin is lower for the distal compartments in all three neuron types, and lowest in Class IV WT.*

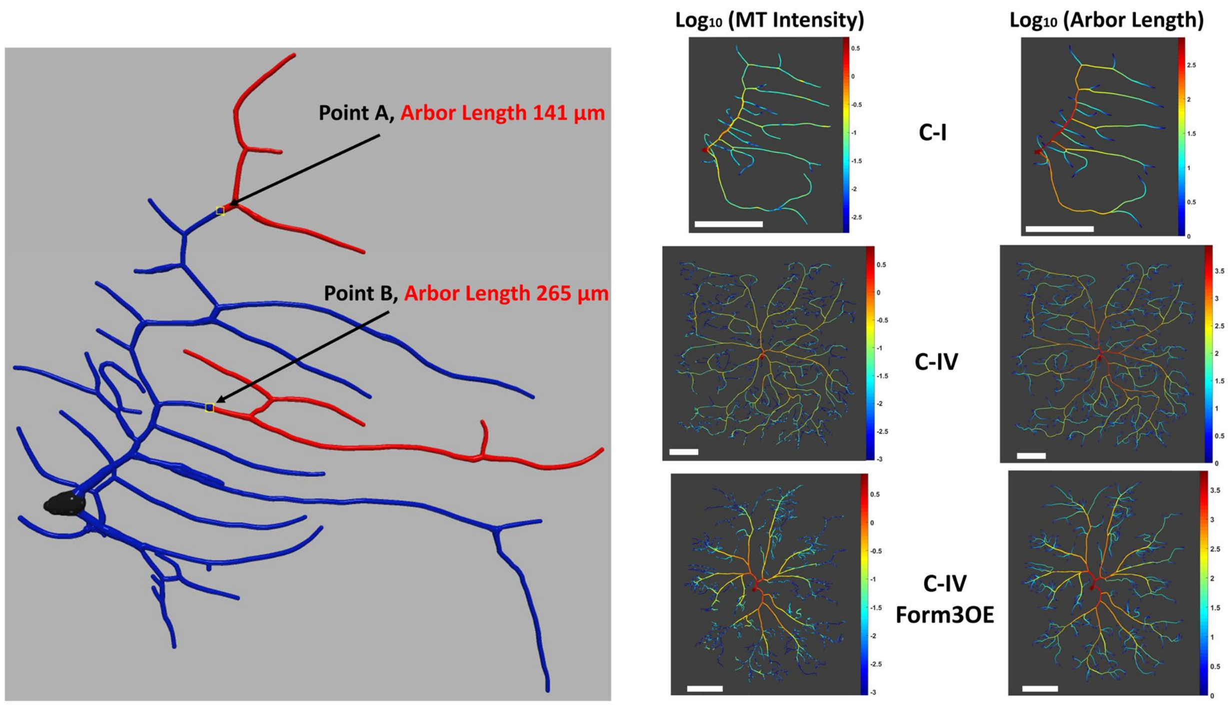


***Figure S4*** *Arbor length of a compartment is defined as the total downstream length from that location. On the left figure, the red dendritic regions are the downstream arbors for the two representative compartments A and B. On the right, microtubule quantity and arbor length for Class I, Class IV and Class IV Form3OE neurons show similar centrifugal decrease.*

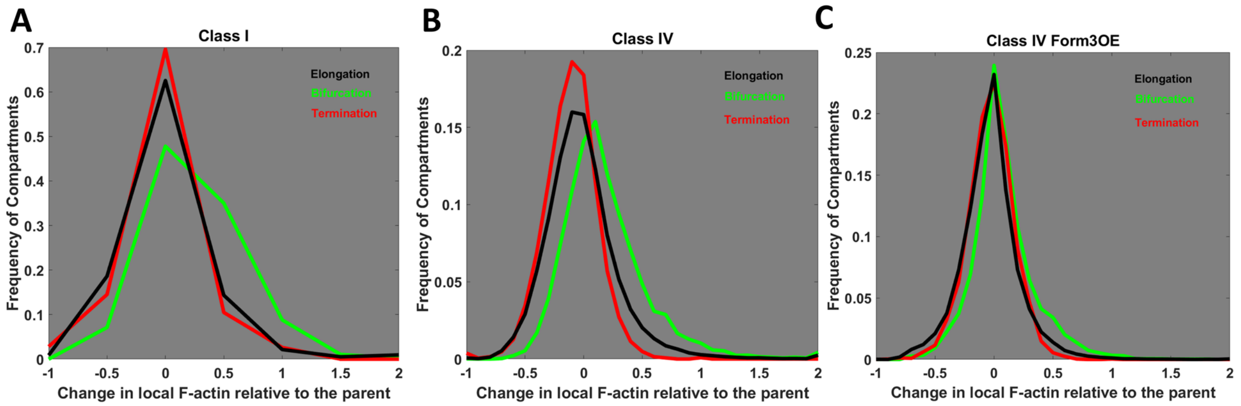


***Figure S5*** *Frequency of the fold change in local F-actin concentration relative to the parent compartment for bifurcating (green), elongating (black) and terminating (red) compartments. A value of 0 indicates unchanged F-actin concentration; a value of -1 indicates complete F-actin disappearance (100% decrease relative to parent compartment); and a value of +1 indicates doubling of F-actin concentration (100% increase relative to parent compartment). Class I (A), Class IV (B) and mutant Class IV Form3OE (C) distributions are shown separately.*

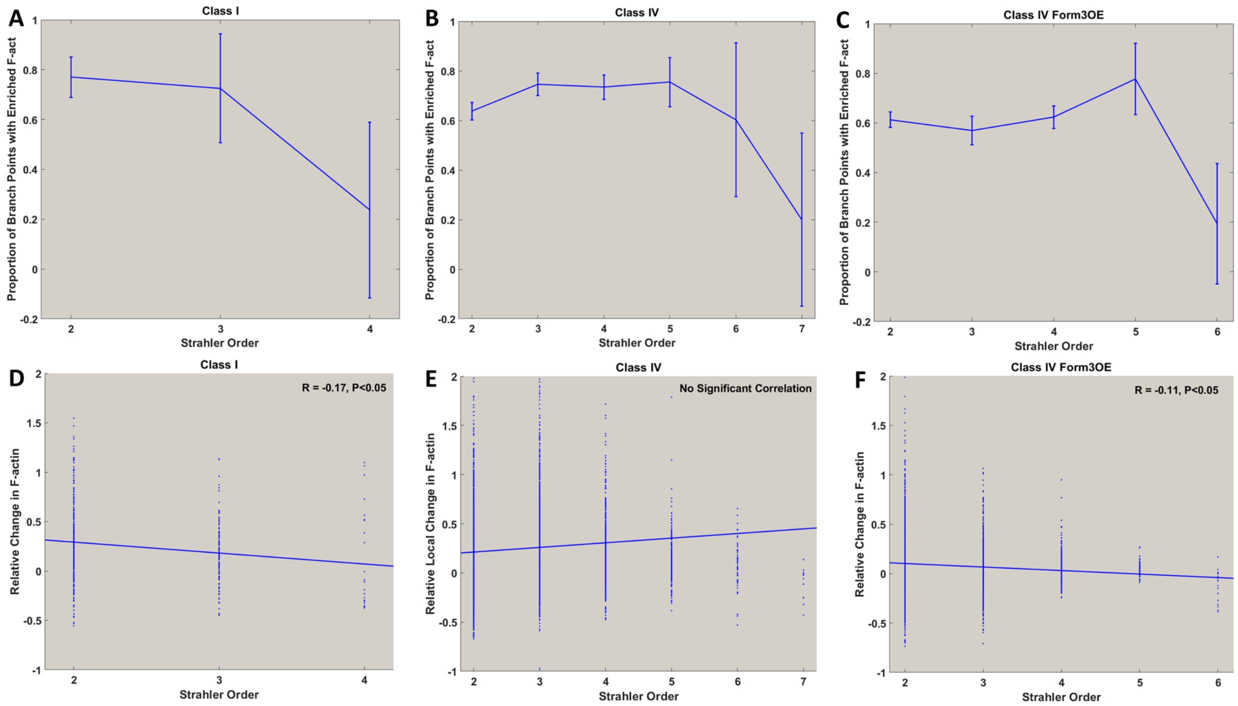


***Figure S6:*** *Strahler order based distribution of branch points and their local change in F-actin . The top row (A, B, C) display the proportion of branch points at each Strahler order with a positive local F-actin change for Class I (A), Class IV WT (B) and Class IV Form3OE (C) neurons. More than half the compartments from all Strahler orders (excluding the somatic regions i.e. the final Strahler order for each neuron type) from all three neuron types were F-actin enriched, i.e. an increase in local F-actin relative to the parent compartment. Error bars display standard deviation. The bottom row (D, E, F) display the correlation between Strahler order and level of F-actin enrichment. Class I (D) and Class IV Form3OE (F) neurons show a negative correlation, indicating a reduction in F-actin enrichment levels while moving away from the terminals and towards the soma. Class IV neurons (E) show no correlation between Strahler order and F-actin enrichment level.*

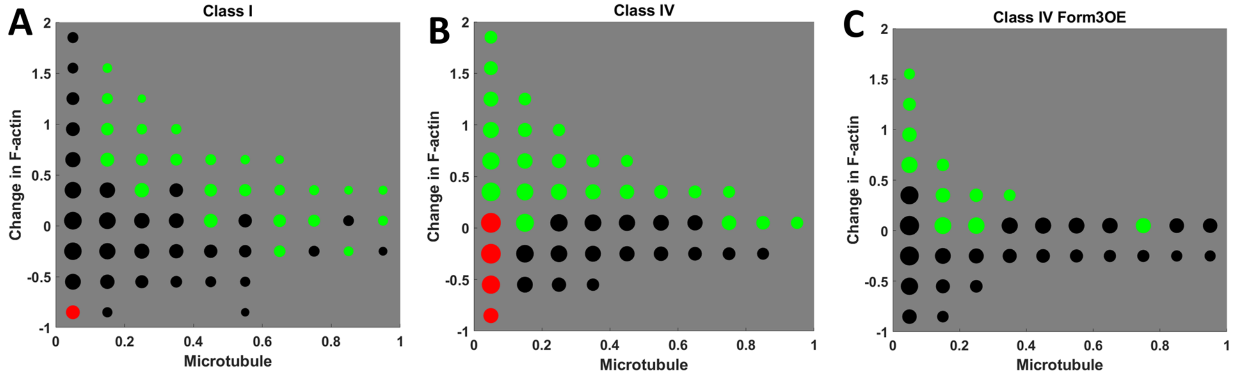


***Figure S7*** *Two-dimensional distribution of all dendritic compartments based on local microtubule and local change in F-actin for Class I (A), Class IV (B) and Class IV Form3OE (C) neurons. The size of each circle is proportional to the log of the number of compartments with MT and F-act quantities in the corresponding range. Green and red circles represent high bifurcation and termination proportions (greater than 8%), respectively. The bifurcating and terminating regions have distinct cytoskeletal composition.*

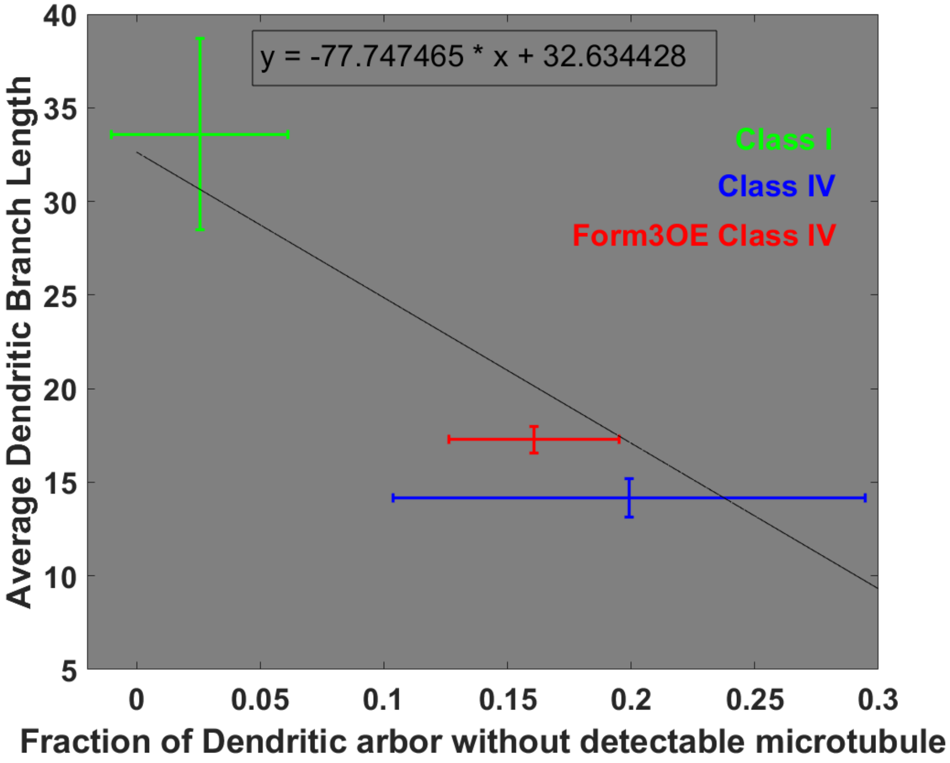


***Figure S8*** *Negative correlation of mean branch length with the proportion of arbor lacking microtubule. The vertical bars represent standard deviation of average branch length and the horizontal bars represent standard deviation of the fraction of arbor lacking microtubule.*

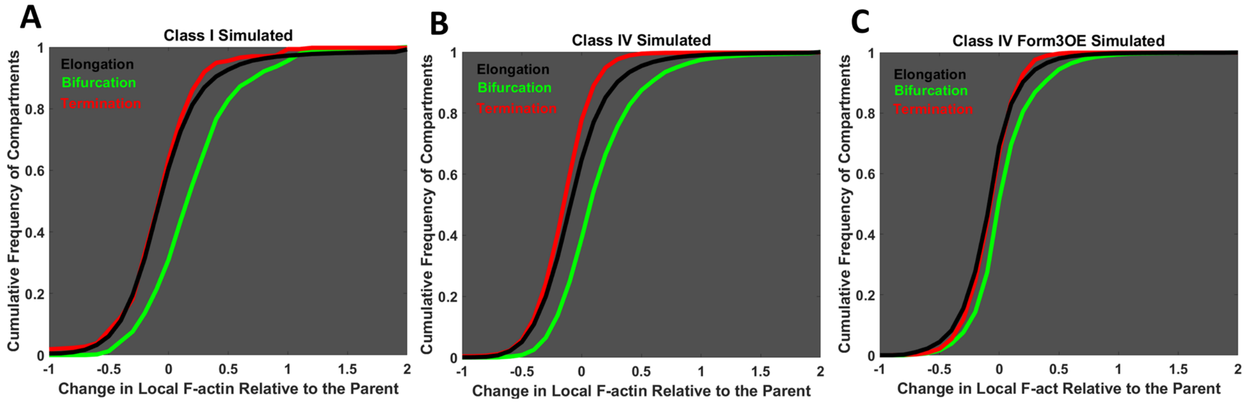


***Figure S9*** *Analysis of simulated neurons. Emergent cumulative frequency of the fold change in local F-actin concentration relative to the parent compartment for bifurcating (green), elongating (black) and terminating (red) compartments. Simulated Class I (A), Class IV (B) and mutant Class IV Form3OE (C) distributions are shown separately. In all three cases, the bifurcating probability distribution function is clearly separate from the terminating and elongating ones, well matching the corresponding experimental measurements.*

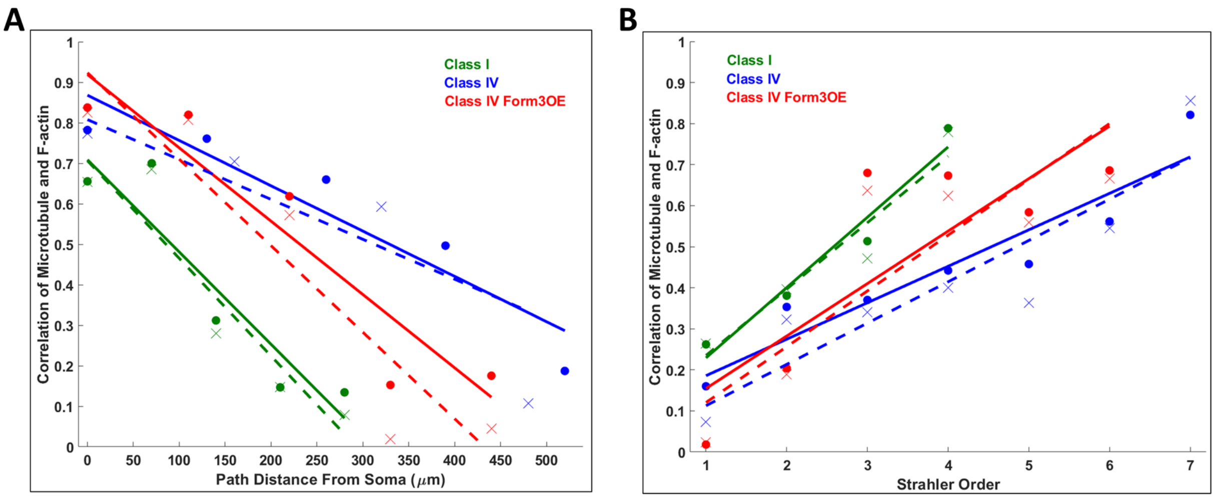


***Figure S10*** *Change of MT-to-F-act correlation as a function of path distance from soma (A), and Strahler order (B). Solid circles represent real neuron correlation distribution and 'x'-marks display the correlation distribution of the simulated neurons. Solid lines represent linear fits for the real neurons, dotted lines represent linear fits for the simulated neurons.*

***Table S1*** *Comparison between real (blue) and simulated (red) neurons of the correlation between microtubule and F-actin expression at different path distances from the soma and of the correlation between microtubule and arbor length in bifurcating and terminating branches.*

| Pearson Correlation Coefficients | Class I | Class IV | Class IV Form3OE |
|---|---|---|---|
| MT-F-act, Closest 20% compartments from the soma (Observed) | 0.98 | 0.88 | 0.89 |
| MT-F-act, Closest 20% compartments from the soma (Emergent) | 0.98 | 0.87 | 0.89 |
| MT-F-act, Farthest 20% compartments from the soma (Observed) | 0.46 | 0.32 | 0.68 |
| MT-F-act, Farthest 20% compartments from the soma (Emergent) | 0.44 | 0.47 | 0.67 |
| MT-Arbor Length bifurcating Branches (Observed) | 0.65 | 0.77 | 0.87 |
| MT-Arbor Length bifurcating Branches (Emergent) | 0.65 | 0.76 | 0.86 |
| MT-Arbor Length terminal Branches (Observed) | 0.26 | 0.16 | 0.02 |
| MT-Arbor Length terminal Branches (Emergent) | 0.26 | 0.07 | 0.02 |

***Table S2*** *Comparison of observed (real neurons) and emergent (simulated neurons) morphological properties.*

| *Properties* | *Class I* | *Class IV* | *Class IV Form3OE* |
|---|---|---|---|
| Total Length (µm) (Observed) | 1680 ± 166 | 19305 ± 1510 | 12591 ± 985 |
| Total Length (µm) (Emergent) | 1653 ± 191 | 19033 ± 1416 | 12154 ± 1761 |
| Total Tips (Observed) | 26 ± 4.8 | 683 ± 44.7 | 365 ± 29.9 |
| Total Tips (Emergent) | 26 ± 5.2 | 673 ± 46.3 | 354 ± 51.5 |
| Length Asymmetry (Observed) | 0.559 ± 0.052 | 0.549 ± 0.017 | 0.570 ± 0.019 |
| Length Asymmetry (Emergent) | 0.559 ± 0.055 | 0.553 ± 0.018 | 0.568 ± 0.021 |
| Topological Asymmetry (Observed) | 0.607 ± 0.077 | 0.610 ± 0.024 | 0.598 ± 0.015 |
| Topological Asymmetry (Emergent) | 0.602 ± 0.089 | 0.608 ± 0.025 | 0.593 ± 0.019 |
| Length Caulescence (Observed) | 0.631 ± 0.107 | 0.450 ± 0.067 | 0.497 ± 0.063 |
| Length Caulescence (Emergent) | 0.616 ± 0.115 | 0.471 ± 0.061 | 0.472 ± 0.080 |
| Topological Caulescence (Observed) | 0.579 ± 0.091 | 0.470 ± 0.080 | 0.488 ± 0.053 |
| Topological Caulescence (Emergent) | 0.565 ± 0.101 | 0.469 ± 0.066 | 0.487 ± 0.033 |

***Table S3*** *Average branch length and ratio between microtubule and F-actin quantities in real and simulated neurons as well as in bifurcating and terminating branches.*

| Neuron Type | C1 | C4 | C4 Form3OE | C1 Sim | C4 Sim | C4 Form3OE Sim |
|---|---|---|---|---|---|---|
| Branch Length (µm) | 33.57 ± 5.11 | 14.17 ± 1 | 17.27 ± 0.7 | 33.45 ± 5.32 | 14.17 ± 1 | 17.21 ± 0.72 |
| MT to F-act ratio | 0.89 ± 0.32 | 0.38 ± 0.14 | 0.25 ± 0.04 | 0.89 ± 0.32 | 0.37 ± 0.14 | 0.26 ± 0.04 |
| Branch Length (Bifurcating) (µm) | 24.91 ± 4.01 | 15.35 ± 1.02 | 16.43 ± 1.22 | 24.75 ± 4.5 | 15.39 ± 1.07 | 16.5 ± 1.21 |
| MT to F-act ratio (Bifurcating) | 1.13 ± 0.27 | 0.55 ± 0.19 | 0.3 ± 0.04 | 1.13 ± 0.26 | 0.55 ± 0.19 | 0.30 ± 0.04 |
| Branch Length (Terminating) (µm) | 41.88 ± 8.07 | 12.98 ± 1.29 | 18.11 ± 0.75 | 41.78 ± 8.07 | 12.96 ± 1.32 | 17.95 ± 0.60 |
| MT to F-act ratio (Terminating) | 0.63 ± 0.39 | 0.07 ± 0.02 | 0.1 ± 0.02 | 0.63 ± 0.39 | 0.08 ± 0.02 | 0.1 ± 0.02 |

***Table S4*** *Pearson correlation coefficients of arbor length against morphological and cytoskeletal parameters at the resolution of single (2 µm long) compartments for simulated neurons.*

| Cell Type | MT quantity | F-act quantity | MT + F-act quantity | Path distance from soma | Branch order |
|---|---|---|---|---|---|
| Class I | **0.66** | 0.54 | 0.61 | -0.47 | -0.34 |
| Class IV | **0.76** | 0.42 | 0.61 | -0.44 | -0.41 |
| Class IV Form3OE | **0.87** | 0.76 | 0.82 | -0.48 | -0.39 |

#### Glossary

**Compartment**: A neuronal compartment in a digital reconstruction is a cylindrical segment whose length is defined by two adjacent points (the current point and its parent point) where the compartment begins at the parent point and ends at the current point. The diameter of the compartment cylinder is the dendritic branch thickness at that location. In this study, the neuronal reconstructions are resampled so as to make each compartment approximately 2 µm long.

**The parent (compartment)** is the compartment immediately before the current compartment. The parent of a given (current or active) compartment is the compartment that is situated immediately before the given compartment when the neuron tree is sorted, i.e. the ordering of compartments within the tree start from the soma and end in terminals. In other words, the parent is the first compartment found while traversing retrogradely from the current compartment towards the soma. All dendritic compartments have one parent compartment. However, a parent compartment may have one to two child compartments. Only terminal compartments (that do not have child compartments) are not parent compartments themselves, and only the root node of a tree has no parent compartment.

**The child (compartment)** is the compartment immediately after the current compartment. The terminal compartments (tips) have no child.

**A bifurcating compartment** is a compartment that branches out into two child compartments.

**An elongating compartment** is a compartment that elongates into one child compartment.

**A terminating compartment** is a compartment that terminates (branch tip), hence has no child compartments.

**Active Compartment:** The term **"active compartment"** is used in the model description. An active compartment is the one which has just been created in simulation, and its cytoskeletal composition is being determined, followed by its topological property, i.e. whether the compartment will elongate further, bifurcate or terminate.

**Path Distance:** The distance (in μm) of a dendritic location from the cell body or the soma is called its path distance.

**Branch Thickness:** The neurite thickness or branch thickness is represented as the diameter of the cylindrical compartments that make up the arbors.

**Integral microtubule:** Descending integral microtubule of a dendritic location is essentially the historical microtubule from the cell body up to that dendritic location.

Descending integral microtubule (***IM***), is mathematically defined for $(i+1)^{th}$ compartment as:

$$\boldsymbol{IM_{i+1} = (IM_i * 0.99) + (MQ_i * 0.01)}$$

where ***MQ*** is the microtubule quantity and the $i^{th}$ compartment is the parent of the $(i+1)^{th}$ compartment. Integral microtubule can equivalently be expressed as:

$$\boldsymbol{IM_{i+1} = 0.99 * MQ_0 + \sum_{j=1}^{i} (0.99)^{i-j} * MQ_j}$$

where $\boldsymbol{MQ_0}$ is the microtubule quantity at the root compartment.

**Arbor length**: Arbor length of a compartment is defined as the total downstream length (of dendritic arbor) from that compartment. Therefore, the arbor length of all dendritic terminals (tips) is zero and the arbor length of the root node of a neuron is that neuron's total neurite length.

**Dendritic length (used as Y-axis in Figure 2A, D and 6A, D):** Dendritic length is the total length of dendrite (in μm) that fall within a topological bin (such as branch order or path distance from soma), averaged across all the neurons of a neurontype. For example, in Fig. 2D, the total length of dendritic branches with the Strahler order of 1 are averaged across all the neurons, and the process is repeated for all the Strahler orders.

**Data Grids:** Three data grids are used in the model. First is the topological event grid from where a topological event (elongation, bifurcation and termination) is sampled. The next two are the bifurcation grid and the elongation grid, from where the cytoskeletal composition of the two bifurcating child compartment or the one elongating child compartment is sampled. A grid in the context of this study, can simply be thought of as a two dimensional matrix, where the two dimensions are local microtubule and local F-actin. Hence, a location within a matrix defines a small range of microtubule and F-actin values. Instead of individual elements, each location within these matrices contain a data bin. Each data bin would get populated when the local microtubule and F-actin quantities measured from the real neurons fall within the microtubule-F-actin range defined by the location of the bin. This is collected from the parent compartments for the two cytoskeletal grids (elongation and bifurcation grids) and from the current compartment for the topological event grid. During the simulation, the cytoskeletal composition of the active compartments is matched with the correct bin, and then data is sampled from the bin to determine the topology of the active compartment, as well as the cytoskeletal composition of its child compartment/compartments.

**Topological event:** The topological event of a compartment is its branching property, i.e. simply the number of child that compartment has. There are three types of topological events (or type of topological compartments), based on whether a compartment is an elongating to a single child compartment or bifurcating to two child compartments or terminating without adding any new child compartment.
